# Disruption of the interfacial membrane leads to *Magnaporthe oryzae* effector re-location and lifestyle switch during rice blast disease

**DOI:** 10.1101/177147

**Authors:** Kiersun Jones, Jie Zhu, Cory B. Jenkinson, Dong Won Kim, Chang Hyun Khang

**Affiliations:** Department of Plant Biology, University of Georgia, Athens, GA 30602

## Abstract

The hemibiotrophic fungus *Magnaporthe oryzae* produces invasive hyphae enclosed in a plant-derived interfacial membrane, known as the extra-invasive hyphal membrane (EIHM), in living rice cells. Little is known about when the EIHM is disrupted and how the disruption contributes to blast disease. Here we show that EIHM disruption correlates with the hyphal growth stage in first-invaded susceptible rice cells. Our approach utilized GFP secreted from invasive hyphae as an EIHM integrity reporter. Secreted-GFP accumulated in the EIHM compartment but appeared in the rice cytoplasm when the EIHM integrity was compromised. Live-cell imaging of secreted-GFP and various fluorescent reporters revealed that EIHM disruption led to rice vacuole rupture and cell death limited to the invaded cell with closed plasmodesmata. We report that EIHM disruption and host cell death are landmarks delineating three distinct infection phases (early biotrophic, late biotrophic, and transient necrotrophic phases) within the first-invaded cell before reestablishment of biotrophy in second-invaded cells. *M. oryzae* effectors exhibited phase-specific localizations, including entry of the apoplastic effector Bas4 into the rice cytoplasm during the late biotrophic phase. Understanding how the phase-specific dynamics are regulated and linked to host susceptibility will offer potential targets that can be exploited to control blast disease.

## INTRODUCTION

Plants grow under constant threat of attack by diverse pathogens, ranging from obligate biotrophs that require living host cells to necrotrophs that benefit from host cell death. Features of both biotrophs and necrotrophs are intricately combined in diverse ways by hemibiotrophic pathogens, which first suppress host cell death during initial biotrophic growth and then induce host cell death during subsequent necrotrophic growth (Horbach et al., 2011). To facilitate infection of the host, pathogens secrete effector proteins in a temporal and spatial manner, some of which modulate host immune responses and cell death, depending on infection stage (Kleemann et al., 2012; Giraldo and Valent, 2013; Toru÷o et al., 2016; Lanver et al., 2017). A major hallmark of biotrophic filamentous pathogens is the specialized intracellular infection structures they produce, such as haustoria and invasive hyphae (IH) (Yi and Valent, 2013; Presti et al., 2015). These structures are separated from the host cytoplasm by a plant-pathogen interface consisting of an apoplastic matrix between the pathogen’s cell wall and a plant-derived membrane (Perfect and Green, 2001; Yi and Valent, 2013). This interface is indispensable for biotrophy and serves a critical role in evading host recognition and acquiring nutrients (Perfect and Green, 2001; Bozkurt et al., 2015).

*Magnaporthe oryzae* is a hemibiotrophic fungus that causes the economically devastating blast disease on rice, wheat and other crops (Khang and Valent, 2010; Cruz and Valent, 2017). For rice alone, the blast fungus destroys harvests that could feed upwards of 60 million people, at a cost of some $66 billion each year (Pennisi, 2010). With the ever increasing demand for food, fundamental understanding of blast infection strategy is more critical than ever to control blast and other related diseases (Khang and Valent, 2010; Pennisi, 2010; Fisher et al., 2012). On the rice leaf surface *M. oryzae* produces a specialized penetration cell, called an appressorium, which produces a narrow penetration peg to breach an epidermal rice cell. *M. oryzae* successively invades living rice cells (biotrophy) before switching to destructive growth associated with macroscopic lesion development and conidiation (necrotrophy) several days after inoculation (Kankanala et al., 2007; Khang and Valent, 2010). Cellular features associated with the early biotrophic invasion have been documented (Figure 1A) (Koga et al., 2004; Kankanala et al., 2007; Mochizuki et al., 2015; Jones et al., 2016b; Shipman et al., 2017). The penetration peg expands to form a filamentous primary hypha, which subsequently differentiates into bulbous IH. The appressorium provides a nucleus to the first bulbous IH cell, which then undergoes multiple cell divisions for 8-12 hours, producing branched IH within the first-invaded host cell before moving into adjacent cells using IH pegs that presumably co-opt plasmodesmata (Kankanala et al., 2007). Both the appressorium and IH undergo a form of semi-closed mitosis, and mitotic nuclei exhibit remarkable constriction and elongation when migrating through the penetration peg or IH pegs (Jones et al., 2016a; Jenkinson et al., 2017). Biotrophic IH are surrounded by a tight-fitting extra-invasive hyphal membrane (EIHM) with irregular localized elaborations (Kankanala et al., 2007; Yi and Valent, 2013). The EIHM appears to originate from the invaginated rice plasma membrane, however it becomes differentiated from it as indicated by differential localization of three rice plasma membrane proteins fused to GFP. GFP:LTi6b continuously outlines IH, OsCERK1:GFP localizes around only the primary hypha, and EL5δ24:GFP generally localizes around the primary hypha but occasionally encases the first bulbous cell (Mentlak et al., 2012; Kouzai et al., 2014; Mochizuki et al., 2015). The endocytotic tracker dye FM4-64 stains rice membranes including the EIHM but is excluded from IH membranes (Kankanala et al., 2007).

**Figure 1.**
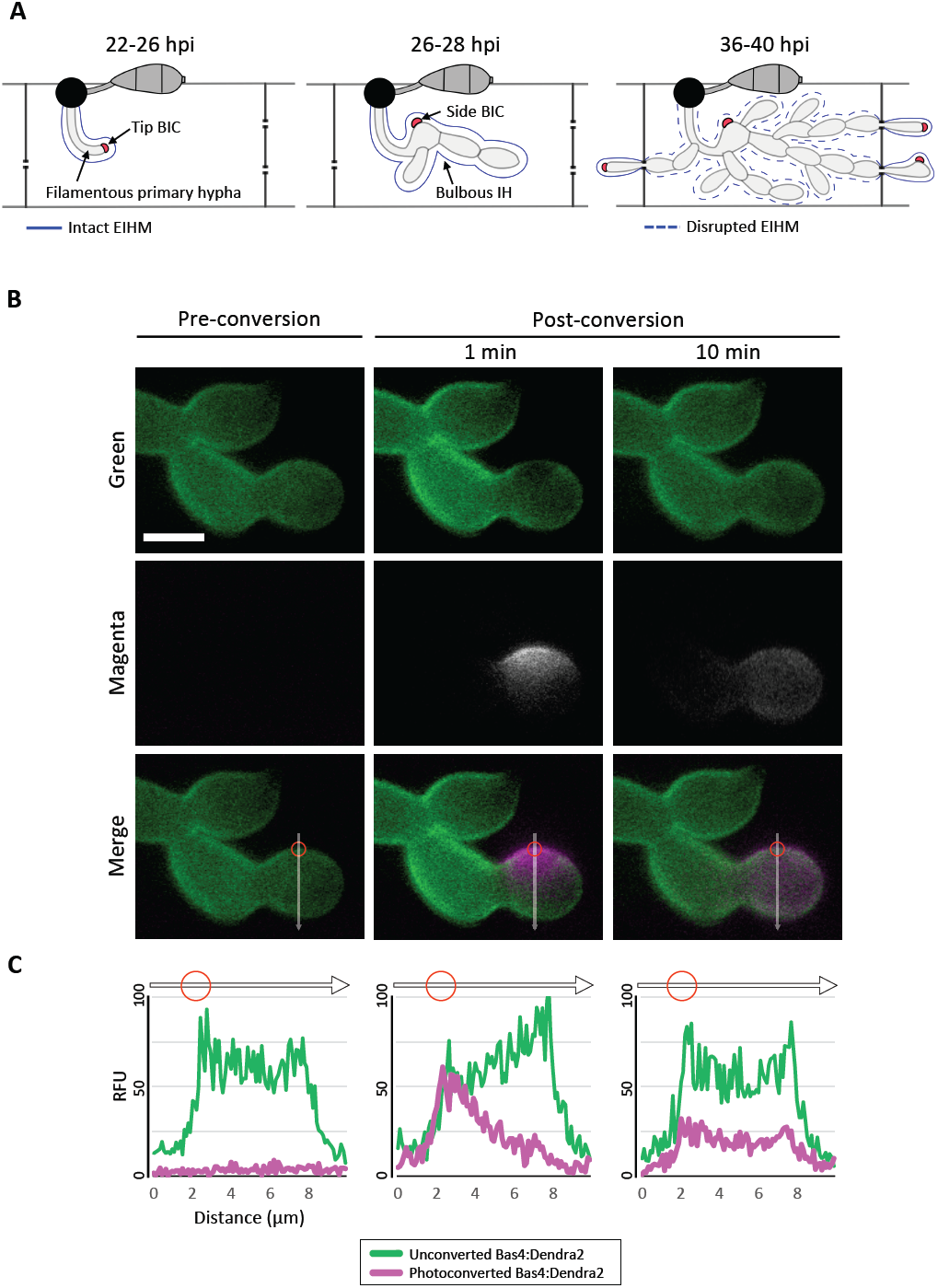
Bas4 is freely-diffusible inside the EIHMx. **(A)** Schematic diagram summarizing the invasion of first and second rice cells by *M. oryzae* IH. At 22-26 hours post inoculation (hpi) a filamentous primary IH grows in the first-invaded host cell where it is surrounded by an intact EIHM. Apoplastic effectors secreted by IH are retained within the EIHMx. In contrast, cytoplasmic effectors enter the host cytoplasm and show preferential accumulation at the tip BIC located at the apex of the primary hypha. At 26-28 hpi the filamentous primary hypha switches to depolarized, asymmetric growth, leaving the BIC subapically associated with the first bulbous IH, becoming a side BIC. Polarized growth resumes from the BIC-associated cell, producing bulbous IH. The EIHM remains intact. At 36-40 hpi the EIHM in first-invaded host cell is disrupted, and IH invade neighboring host cells. Every IH that invades an adjacent rice cell is surrounded by a new EIHM and associated with a new BIC. **(B)** IH of *M. oryzae* strain CKF1737 expressing EIHMx-localized effector Bas4 fused to the green-to-red photoconvertible fluorescent protein Dendra2, invading a rice cell at 29 hpi. Shown are single plane confocal images of separate fluorescence (top and middle panels) and merged fluorescence (bottom panels). Left: Before photoconversion, green Bas4:Dendra2 fluorescence localized throughout the EIHMx. Middle: One minute after selective photoconversion (region indicated by the red circle), red Bas4:Dendra2 fluorescence (magenta and pseudo-colored white) diffused into the surrounding EIHMx. Right: Ten minutes after photoconversion, the red Bas4:Dendra2 further diffused. White arrows indicate the locations of fluorescence intensity linescans shown in **(C)**. Bar = 5 μm. **(C)** Linescans showing the relative fluorescence intensity between unconverted Bas4:Dendra2 (green) and photoconverted Bas4:Dendra2 (magenta), corresponding to the location of the white lines in **(B)**. Red circles show the photoconverted region in **(B)**. Units are relative fluorescence units (RFU; y-axis) and distance in μm (x-axis).

Although the detailed structures of interfacial membranes vary considerably depending on the specific plant-pathogen interaction, the EIHM formed during *M. oryzae* invasion of rice appears analogous to the extrahaustorial membrane (EHM) that encases haustoria of obligate biotrophs; separating the pathogen from the host cytoplasm (Kankanala et al., 2007; Yi and Valent, 2013). However, a key difference exists due to the deviating growth styles of obligate biotrophs and hemibiotrophs. Haustoria are terminal structures, thus the EHM is constructed over a finite surface area, and integrity must be maintained to facilitate obligate biotrophy. In contrast, *M. oryzae* and other hemibiotrophs produce IH that continue to grow and spread into adjacent cells, therefore EIHM construction and integrity maintenance are likely to be perturbed as host cells become increasingly stressed, and membrane materials become exhausted. In fact, several studies noted that the EIHM could lose integrity during IH growth in first-invaded rice cells (Mosquera et al., 2009; Khang et al., 2010; Mochizuki et al., 2015). In addition to the loss of EIHM integrity, first-invaded cells exhibit shrinkage and rupture of the central vacuole as well as loss of rice cell viability around the time IH move into adjacent living rice cells (Kankanala et al., 2007; Mochizuki et al., 2015; Jones et al., 2016b).

Live-cell imaging of *M. oryzae* strains expressing fluorescently-tagged effector proteins have shown differential subcellular localization in the host cell after they are secreted from IH via two distinct protein secretion pathways (Giraldo et al., 2013). Apoplastic effectors (i.e., Bas4, Bas113, and Slp1) are retained in the extrainvasive hyphal matrix (EIHMx), which is the sealed apoplastic compartment formed between the EIHM and IH cell wall (Kankanala et al., 2007; Mosquera et al., 2009; Mentlak et al., 2012; Yi and Valent, 2013). In contrast, cytoplasmic effectors (i.e., Pwl2, Bas1, Bas107, and AvrPiz-t) preferentially accumulate in the biotrophic interfacial complex (BIC), which is a plant-derived localized structure (Khang et al., 2010; Park et al., 2012; Giraldo et al., 2013). Increasing evidence supports that BICs are the site of effector translocation into the host cytoplasm (Khang et al., 2010; Giraldo et al., 2013; Giraldo and Valent, 2013). The two-stage development of BICs, from tip-to side-BIC, occurs in conjunction with IH differentiation from filamentous to bulbous in successfully invaded living rice cells (Figure 1A) (Khang et al., 2010; Shipman et al., 2017). In the first-invaded cell the tip BIC appears at the apex of the filamentous primary hypha where it remains throughout the polarized hyphal growth. When the primary hypha switches to depolarized growth, the BIC is left behind at a subapical position on the first bulbous IH cell. Polarized growth is subsequently resumed producing bulbous IH that branch to colonize the first host cell. Only a single BIC is present in each first-invaded cell, whereas multiple BICs can be present in subsequently invaded adjacent cells, one associated with each IH entering them (Figure 1A). Cell-to-cell movement of effectors during the early infection stage and of IH at the later infection stage indicates symplastic continuity is maintained via open plasmodesmata between the first-invaded cell and uninvaded adjacent cells (Kankanala et al., 2007; Khang et al., 2010). However, live-cell imaging of *M. oryzae*-invaded rice cells coupled with fluorescein diacetate (FDA) staining provided evidence that plasmodesmata permeability was reduced during host vacuole shrinkage and after vacuole rupture in the first-invaded cell (Jones et al., 2016b). Despite these advances in our understanding of rice blast infection dynamics, little is known as to how these cellular dynamics are coordinated.

In this study, we used live-cell imaging of susceptible rice cells invaded by *M. oryzae* transformants expressing various fluorescent reporters to investigate infection development in first-and second-invaded cells. We show that EIHM disruption occurred in the first-invaded cell, contingent on IH growth stage, followed by shrinkage and eventual rupture of the rice vacuole before IH spread into adjacent cells. Vacuole rupture coincided with host cell death, which occurred in a contained manner with presumed closure of plasmodesmata. We demonstrate that *M. oryzae* undergoes three distinct infection phases in the first-invaded cell before reestablishing biotrophy in the second-invaded cells. *M. oryzae* effectors exhibited phase-specific localization. Understanding how the phase-specific cellular dynamics are regulated and linked to host susceptibility will offer potential targets that we can exploit to control blast disease.

## RESULTS

### Secreted proteins are mobile in the EIHMx

To investigate the nature of the rice-*M. oryzae* interface, we generated an *M. oryzae* transformant expressing Bas4 as a translational fusion to the photoconvertible fluorescent protein Dendra2, which can be irreversibly changed from green to red fluorescence upon irradiation with UV light (Gurskaya et al., 2006). Bas4 is an *M. oryzae* effector protein that contains a N-terminal signal peptide (SP; 21 amino acids) which mediates the secretion of the leaderless Bas4 (81 amino acids) into the EIHMx (Mosquera et al., 2009; Khang et al., 2010; Giraldo et al., 2013). During invasion of rice cells the transformant showed bright green fluorescence around IH, indicating that Bas4:Dendra2 was indeed secreted into the EIHMx. This localization pattern was consistent with patterns that have been observed for Bas4 fused to other fluorescent proteins, such as EGFP or mCherry (Mosquera et al., 2009; Khang et al., 2010; Mochizuki et al., 2015). To investigate the mobility of Bas4:Dendra2 in the EIHMx, we selectively photoconverted a small region of Bas4:Dendra2 and then monitored dynamics of both the converted red and the unconverted green fluorescence (Figure 1B). Photoconverted Bas4:Dendra2 progressively diffused into the surrounding EIHMx over the next several minutes, meanwhile unconverted Bas4:Dendra2 diffused into the photoconverted region (Figure 1B and 1C). These results indicated that secreted proteins are diffusible in the EIHMx.

### Secreted GFP as a reporter for EIHM integrity and other host cellular dynamics

Considering the mobility of secreted Bas4:Dendra2 within the EIHMx (Figure 1B), we reasoned that secreted GFP (sec-GFP) could be used to monitor the integrity of the EIHM. In the case of a completely intact EIHM, sec-GFP would be retained exclusively within the EIHMx; conversely, if EIHM integrity is compromised, sec-GFP would spill from the EIHMx into the rice cell lumen. To test this, we used an *M. oryzae* strain expressing a fusion of GFP with the Bas4 signal peptide coupled with the lipophilic dye FM4-64 staining. FM4-64 was previously shown to label fungal membranes, notably at septa, only when the EIHM integrity was compromised (Kankanala et al., 2007). Consistent with this, we found that FM4-64 was visible at fungal septa only when sec-GFP appeared in the rice cell lumen (see Supplemental Figure 1 online). These results demonstrated the utility of sec-GFP localization to reveal EIHM integrity.

Three distinct patterns of sec-GFP localization were identified by live-cell imaging of first-invaded rice cells infected with *M. oryzae* strain CKF1996 expressing sec-GFP together with cytoplasmic tdTomato (n > 100). The first pattern was sec-GFP exclusively localized in the EIHMx, indicating an intact EIHM (Figure 2A; infection 1). The second pattern was sec-GFP localized in the rice cytoplasm excluded from the shrunken vacuole (Figure 2A; infection 2; Supplemental Figure 2 online). The third pattern was homogenous distribution of sec-GFP throughout the rice cell with the collapsed vacuole (Figure 2A; infection 3 and 4). These results were consistent with recently reported shrinkage and collapse of the central vacuole in *M. oryzae*-invaded rice cells (Mochizuki et al., 2015; Jones et al., 2016b). Interestingly, spilled sec-GFP did not diffuse into neighboring rice cells (Figure 2; see Supplemental Figures 1 and 2 online), suggesting plasmodesmata were closed. Together, these results showed that sec-GFP localization provides a robust assay for the integrity of the EIHM as well as the state of the host vacuole and plasmodesmata permeability after EIHM disruption.

**Figure 2.**
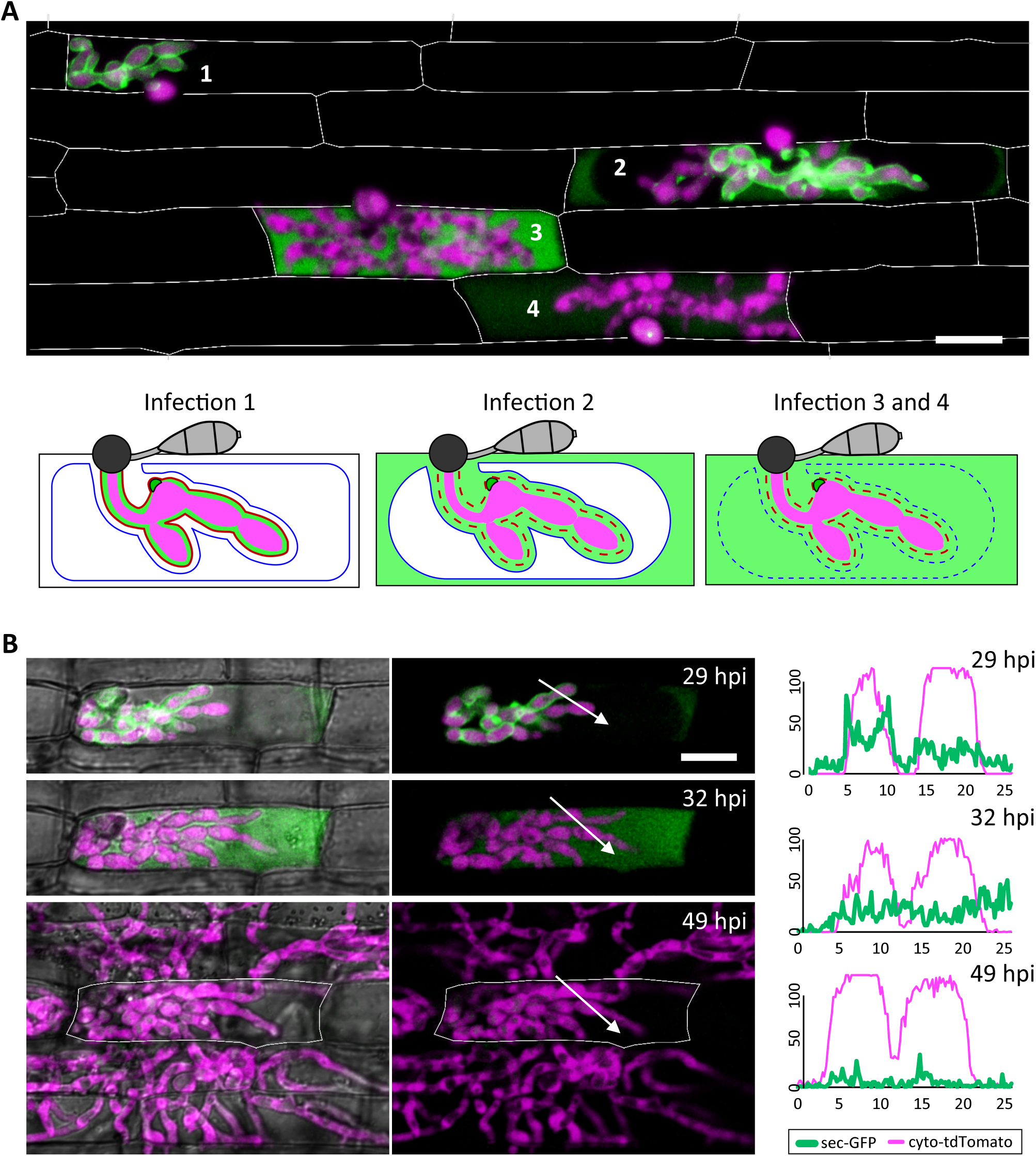
The EIHM loses integrity during invasion of the first host cell. **(A)** A merged fluorescence projection of *M. oryzae* strain CKF1996, expressing sec-GFP (green) and cytoplasmic tdTomato (magenta) in first-invaded rice cells at 32 hpi. Infections are representative of the patterns observed for sec-GFP localization: retention within the EIHMx (infection 1), spilled into the rice cytoplasm with exclusion from the vacuole (infection 2), and spilled homogenously into the rice cell lumen with a ruptured vacuole (infection 3 and 4). Rice cell walls are denoted by white outlines. The same three sec-GFP patterns are schematically illustrated with the addition of the EIHM membrane (red line) and the vacuole membrane (blue line). Disrupted membranes are indicated by a dotted line. Bar = 20 μm. **(B)** Shown are merged fluorescence and bright-field (left) and merged fluorescence alone (right) of a time-lapsed CKF1996 infection from 29 to 49 hpi. At 29 hpi, sec-GFP fluorescence (green) was partially spilled from the EIHMx into the rice cytoplasm and excluded from the vacuole. Three hours later, all sec-GFP fluorescence was homogenously distributed throughout the rice cell. After another 17 hours, viable IH (magenta) had exited the first-invaded host cell (white outline) and had successfully invaded and colonized multiple adjacent rice cells. White arrows denote the locations used for generating the fluorescence intensity linescans. Bar = 20 μm. The linescans measure the relative fluorescence intensity of sec-GFP (green) and cytoplasmic tdTomato (magenta). At 29 hpi, two neighboring hyphae have different localization patterns of sec-GFP; one hypha is outlined by sec-GFP fluorescence (represented by the two green fluorescence peaks) while the other hypha does not show sec-GFP outlining. At 32 hpi, the same two hyphae both now lacked sec-GFP outlining. At 49 hpi, the same IH did not show significant sec-GFP fluorescence. Units are relative fluorescence units (RFU; y-axis) and distance in μm (x-axis).

### Fungal colonization continues after EIHM disruption

Not all infections result in successful colonization of the host even when it is a susceptible interaction (Heath et al., 1990). We therefore considered that EIHM disruption could be associated with failed infection. Thus, we tested if IH growth becomes arrested after EIHM disruption in the first-invaded rice cell. Time-lapse imaging of rice cells invaded by the *M. oryzae* strain CKF1996 (sec-GFP and cytoplasmic tdTomato) showed that IH continued to colonize host cells after the EIHM was disrupted during invasion of the first host cell (Figure 2B; n=15). The growth of IH in the time-lapsed infections was consistent with freshly prepared control infections that were not subject to any potential imaging-related stress (data not shown). Therefore, we concluded that disruption of the EIHM during invasion of the first cell is characteristic of successful colonization of rice by *M. oryzae.* These time-lapse imaging results also revealed that sec-GFP first spilled into the rice cytoplasm with exclusion from the shrinking central vacuole, followed by homogenous distribution throughout the rice cell upon vacuole rupture and that IH subsequently spread into adjacent rice cells (Figure 2B).

### EIHM disruption process

To gain insight into how the EIHM is disrupted, we observed the early stage of sec-GFP spilling in first-invaded cells (n=18) and identified four features associated with EIHM disruption: (1) the initial loss of sec-GFP from the EIHMx appeared to occur near the tips of growing IH (Figure 3; asterisks, linescan a^0^ and a), indicating the initial EIHM disruption at the expanding hyphal region, (2) sec-GFP outline disappeared from some IH while others retained sec-GFP outline (Figure 3A; increase in the number of IH denoted by white asterisks), suggesting that EIHM disruption occurred separately at different locations and was not globally executed, (3) IH that had lost sec-GFP outline did not recover outline, and IH continued to grow without accumulation of new sec-GFP outlining (Figure 3; linescans b^0^, b, and c), indicating that disruption of the EIHM was permanent, and (4) sec-GFP frequently appeared in a punctate pattern at the surface of IH associated with the loss of sec-GFP outline (Figure 3; white arrowheads, linescan d). These puncta varied significantly in terms of quantity, intensity, and duration. The nature of these sec-GFP puncta remains to be determined.

**Figure 3.**
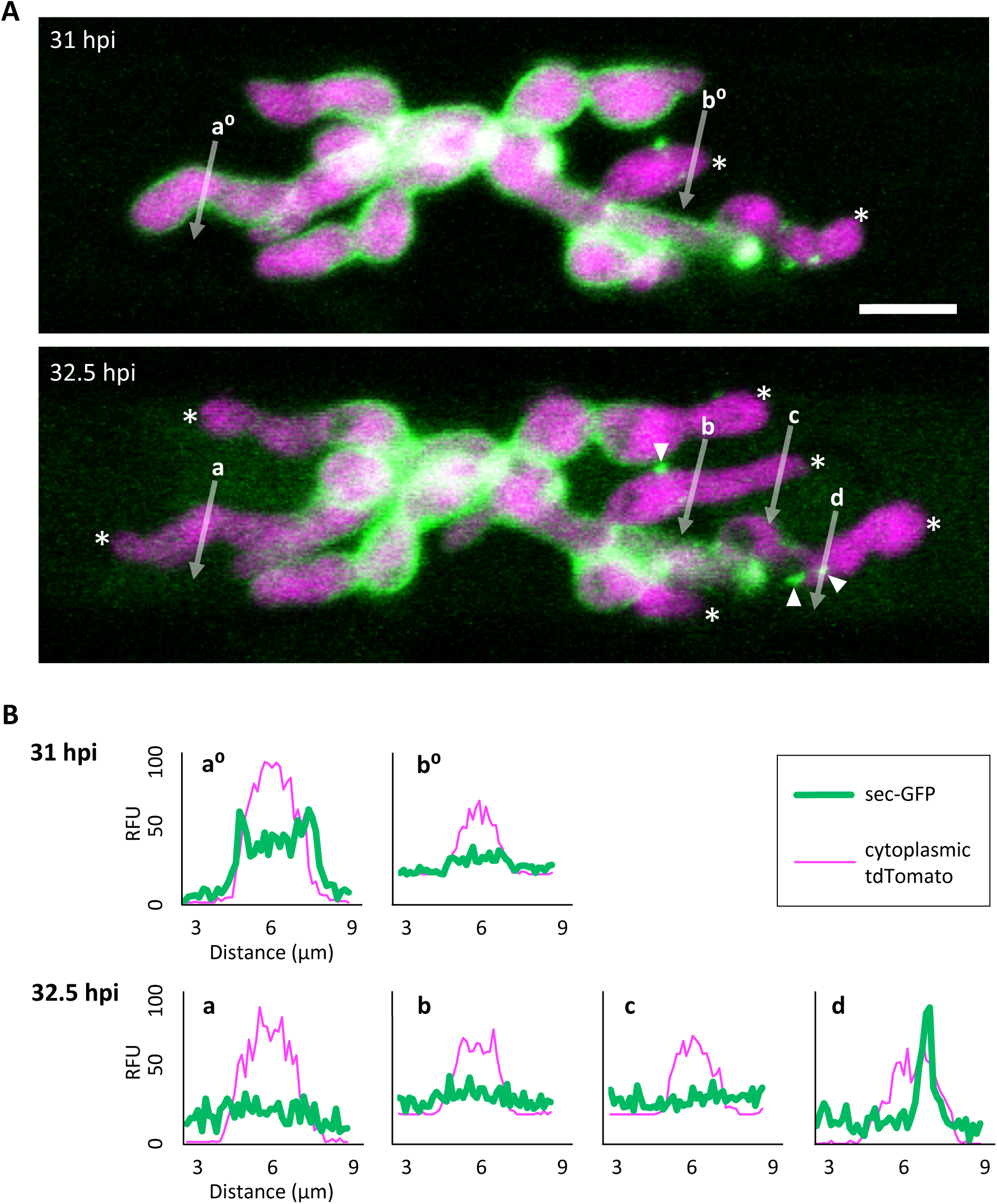
Sec-GFP localization changes during the process of EIHM disruption. **(A)** *M. oryzae* CKF1996 expressing sec-GFP (green) and cytoplasmic tdTomato (magenta) invading a rice cell. Shown are merged fluorescence projections of informative focal planes from the same infection site at 31 (top) and 32.5 hpi (bottom). At 31 hpi sec-GFP localized in the host cytoplasm (not shown in the field of view) and outlined most IH (IH without asterisks and linescan a^0^). Two IH had lost sec-GFP outlining (white asterisks and linescan b^0^). The same infection 1.5 hours later had increased sec-GFP accumulation in the host cell (now visible in the field of view) and loss of sec-GFP outline from additional IH (increase from 2 to 6 white asterisks). White letters and arrows denote the locations used to generate the fluorescence intensity linescans shown in **(B)**. White arrowheads indicate sec-GFP puncta associated with disrupted EIHM. Bar = 10 μm. **(B)** Linescans showing the relative fluorescence intensities (RFU; y-axis, distance in μm; x-axes) of sec-GFP (green) and cytoplasmic tdTomato (magenta) from **(A),** highlighting features of sec-GFP fluorescence localization changes during the process of EIHM disruption. Linescans at 31 hpi show an IH with a sec-GFP outline (a^0^) and an IH without a sec-GFP outline (b^0^). Linescans generated from the same locations at 32.5 hpi show new loss of sec-GFP outline (a) and maintained absence of sec-GFP outlining after initial loss (b). Additional linescans show: new IH growth after sec-GFP outline loss without accumulation of new sec-GFP outline (c), and a sec-GFP puncta associated with outline loss (d). Note that linescans generated at 32.5 hpi clearly show sec-GFP fluorescence spilled into the rice cell lumen (green fluorescence not associated with IH).

### The occurrence of EIHM disruption increases with IH growth stage

We determined the relationship between EIHM disruption and IH growth stage by analyzing 390 rice cells infected with *M. oryzae* strain CKF2187, expressing sec-GFP and a translational fusion of tdTomato to histone H1 (H1:tdTomato; Figure 4). The growth stage was determined by counting H1:tdTomato-tagged nuclei because each IH cell contains one nucleus. We implemented an empirically-derived image analysis method to increase the sensitivity to low intensity sec-GFP fluorescence in the host cytoplasm so that infections at the early stage of sec-GFP spill were correctly scored (see Supplemental Figure 3 online). We found that of 390 infections, 235 had an intact EIHM, and 155 had a disrupted EIHM (Figure 4D). Most of the 155 infections with a disrupted EIHM showed sec-GFP spilled into either the rice cytoplasm or homogenously with the disrupted vacuole. However, about ∼7% (n=11 of 155) showed alternative patterns of localization, such as brighter accumulation of sec-GFP in the vacuole (see Supplemental Figure 4 online). We found that EIHM disruption occurred as early as the 3 to 4 nuclear stage, although at a low frequency (Figure 4D; n=3 of 38). By the 13-14 nuclear stage EIHM disruption frequency reached about 50% (Figure 4D; n=17 out of 33). All infections at the 21 or more nuclear stage exhibited EIHM disruption (Figure 4D; n=21 of 21). These results showed that the occurrence of EIHM disruption in the first-invaded cell increased proportionally with IH growth.

**Figure 4.**
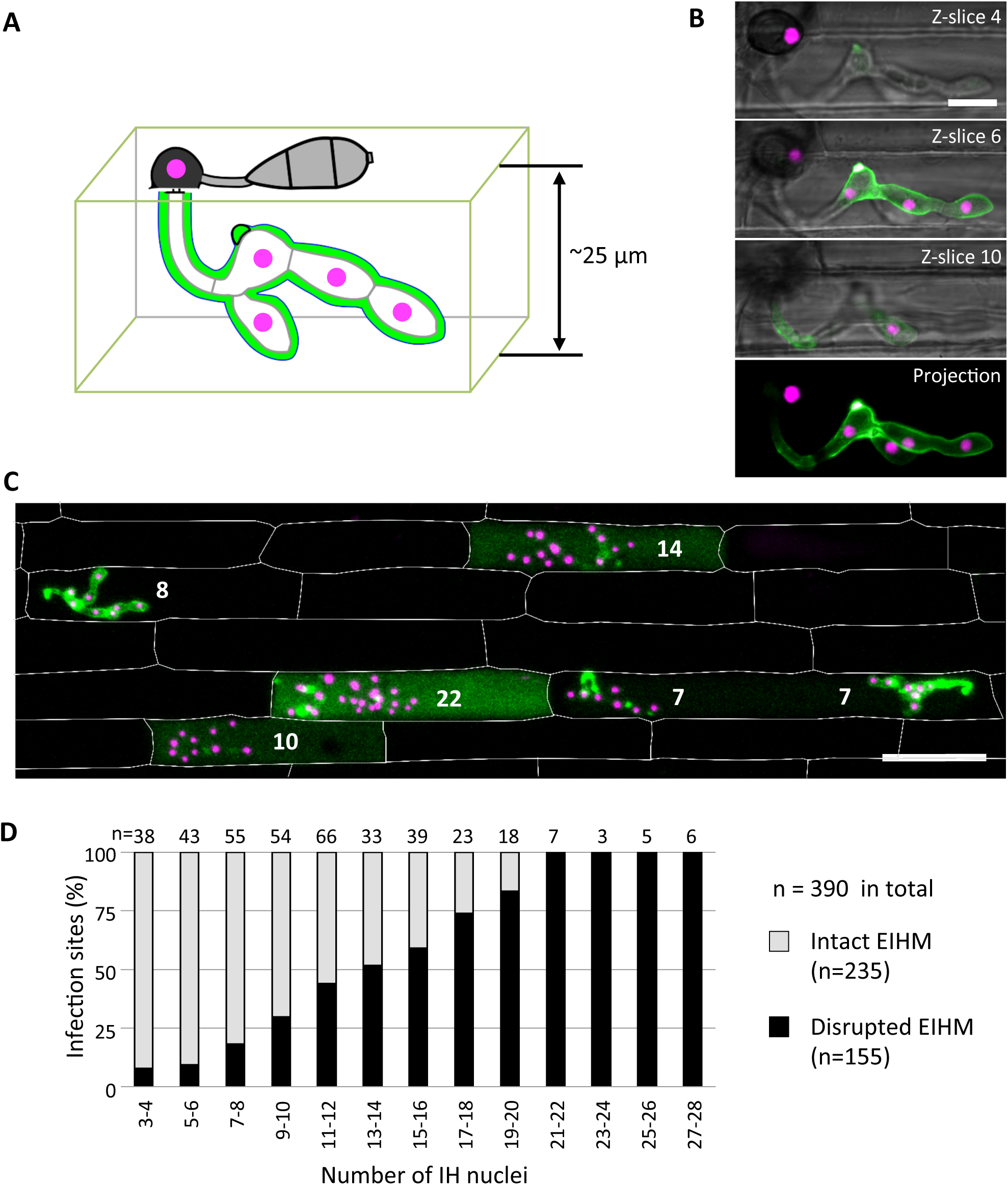
The occurrence of EIHM disruption increases proportionally with nuclear stage. **(A)** Schematic diagram three-dimensionally depicting an epidermal rice cell invaded by *M. oryzae* expressing sec-GFP (green) and nuclear tdTomato (magenta). **(B)** and **(C)** Confocal images of rice cells invaded by *M. oryzae* CKF2187 expressing sec-GFP (green) and nuclear tdTomato (magenta). **(B)** Images of the same infection taken at different focal planes. The top three panels show merged fluorescence and bright-field from individual focal planes (indicated in the top right corner), while the last panel shows a merged fluorescence projection of all focal planes; 12 z-slices in total spanning 24 μm over the z-axis. Note that all five fungal nuclei are fully visible only in the projection view. Bar = 10 μm. **(C)** A merged fluorescence projection of infected rice cells at 30 hpi showing different patterns of sec-GFP localization at different IH growth stages determined by the nuclear number for each infection (white numbers). The infection to the far left shows EIHMx-localized sec-GFP (intact EIHM), while the other infections show host-localized sec-GFP (disrupted EIHM). Note the infected rice cell at the bottom right was invaded by two separate appressoria. Rice cell walls are indicated by white outlines. Bar = 50 μm. **(D)** A plot of the frequency of EIHM disruption according to fungal nuclear number for 390 infections of *M. oryzae* CKF2187 in the first-invaded host cell between 28 and 33 hpi. Sec-GFP localization patterns revealed the distribution of intact (n=235) and disrupted (n=155) EIHMs (y-axis) at each group of two nuclear stages (x-axis).

### Death of the first-invaded rice cell coincides with vacuole rupture

To determine when viability of the first-invaded cell is lost, we infected rice cells with *M. oryzae* strain CKF315 expressing sec-GFP and then stained with propidium iodide (PI) just before microscopy. PI staining was previously used to identify dead rice cells during blast invasion because it indiscriminately labels plant cell walls but only labels nuclei when the plasma membrane is permeabilized, thereby indicating a dying or dead cell (Jones et al., 2016b). We found that infections with an intact EIHM did not show nuclear PI labelling, indicating invaded rice cells were viable as expected (Figure 5A and 5B; ‘a’). Infected cells with a disrupted EIHM and an intact vacuole were typically viable (Figure 5A and 5B; ‘b’), however, some were dead based on nuclear PI labelling (Figure 5A and 5B; ‘c’). Conversely, infected cells with a disrupted EIHM and a ruptured vacuole were rarely viable (Figure 5A and 5B; ‘d’) with the majority appearing dead (Figure 5A and 5B; ‘e’). These results indicated that before IH spread into adjacent rice cells, the first-invaded rice cell died nearly concurrent with vacuole rupture. The association between vacuole rupture and cell death was further confirmed by time-lapse imaging (Figure 5C and 5D). Uninvaded rice cells that were adjacent to a dead first-invaded cell remained viable (Figure 5A; ‘e’).

**Figure 5.**
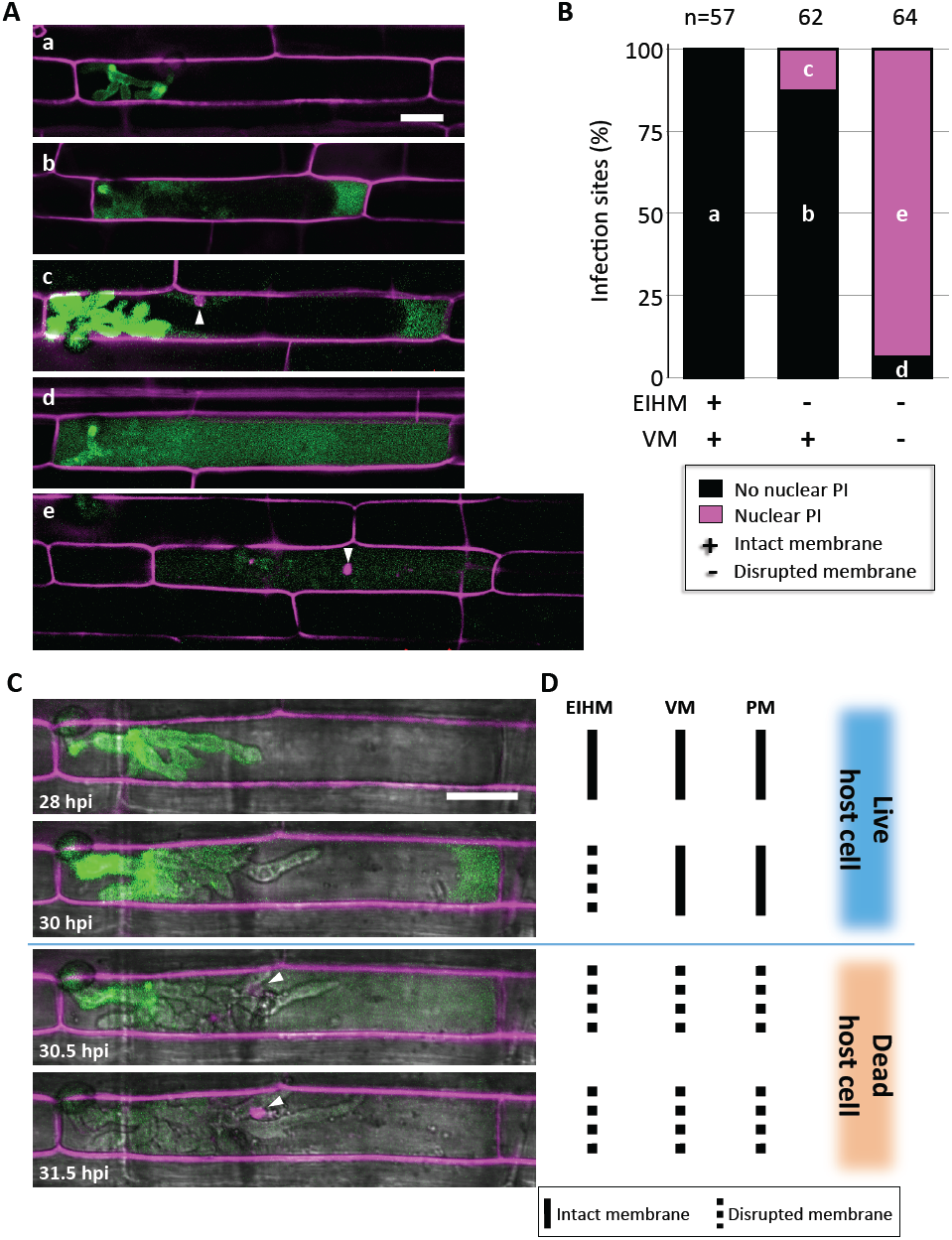
Vacuole rupture indicates host cell death. **(A)** and **(C)** Rice cells invaded by *M. oryzae* CKF315 (sec-GFP; green) and stained with propidium iodide (PI; magenta). Shown are single plane confocal images of merged fluorescence **(A)** and merged bright-field and fluorescence **(C)**. White arrowheads indicate rice nuclei stained with PI. Bars = 20 μm. **(A)** Representative images from the 183 infections from 29 to 34 hpi showing the five sec-GFP and PI localization patterns observed: (a) EIHMx-exclusive sec-GFP without nuclear PI stain, (b) sec-GFP spilled into the rice cytoplasm without nuclear PI stain (c) same as (b) but with nuclear PI stain, (d) sec-GFP homogenized throughout the host cell lumen without nuclear PI stain, (e) same as (c) but with nuclear PI stain. **(B)** Graph showing the distribution of the five sec-GFP and PI fluorescence patterns shown in **(A)** for all 183 infections. Black bars and magenta bars represent infections without nuclear PI staining and with nuclear PI staining, respectively. EIHM = extrainvasive hyphal membrane. VM = host vacuole membrane. **(C)** Time-lapse series of a PI-stained CKF2180 infection from 28 to 31 hpi showing the typical progression of sec-GFP and PI fluorescence localization changes, consistent with quantitative results in **(B)**. Note that appearance of host nuclear PI fluorescence coincided with homogenization of host-localized sec-GFP fluorescence. **(D)** Schematic diagram summarizing the states of host membranes, corresponding to the infection shown in **(C)**. EIHM = extrainvasive hyphal membrane. VM = host vacuole membrane. PM = host plasma membrane.

### IH undergo transient necrotrophic-like growth in the dead first-invaded host cell

Using sec-GFP and H1:tdTomato reporters to correlate the timing of vacuole rupture and IH growth stage in the first-invaded cell, we found that the vacuole ruptured at an average nuclear stage of 28 (Figure 6B; gray bars, range 22-34). The average then increased by six (Figure 6B; black bars; range 1-12) to an average nuclear stage of 34 when IH began to invade adjacent rice cells. The time that elapsed between rupture of the vacuole and the spread of IH into adjacent cells was 1.3 to 2.5 hours (Figure 6B).

**Figure 6.**
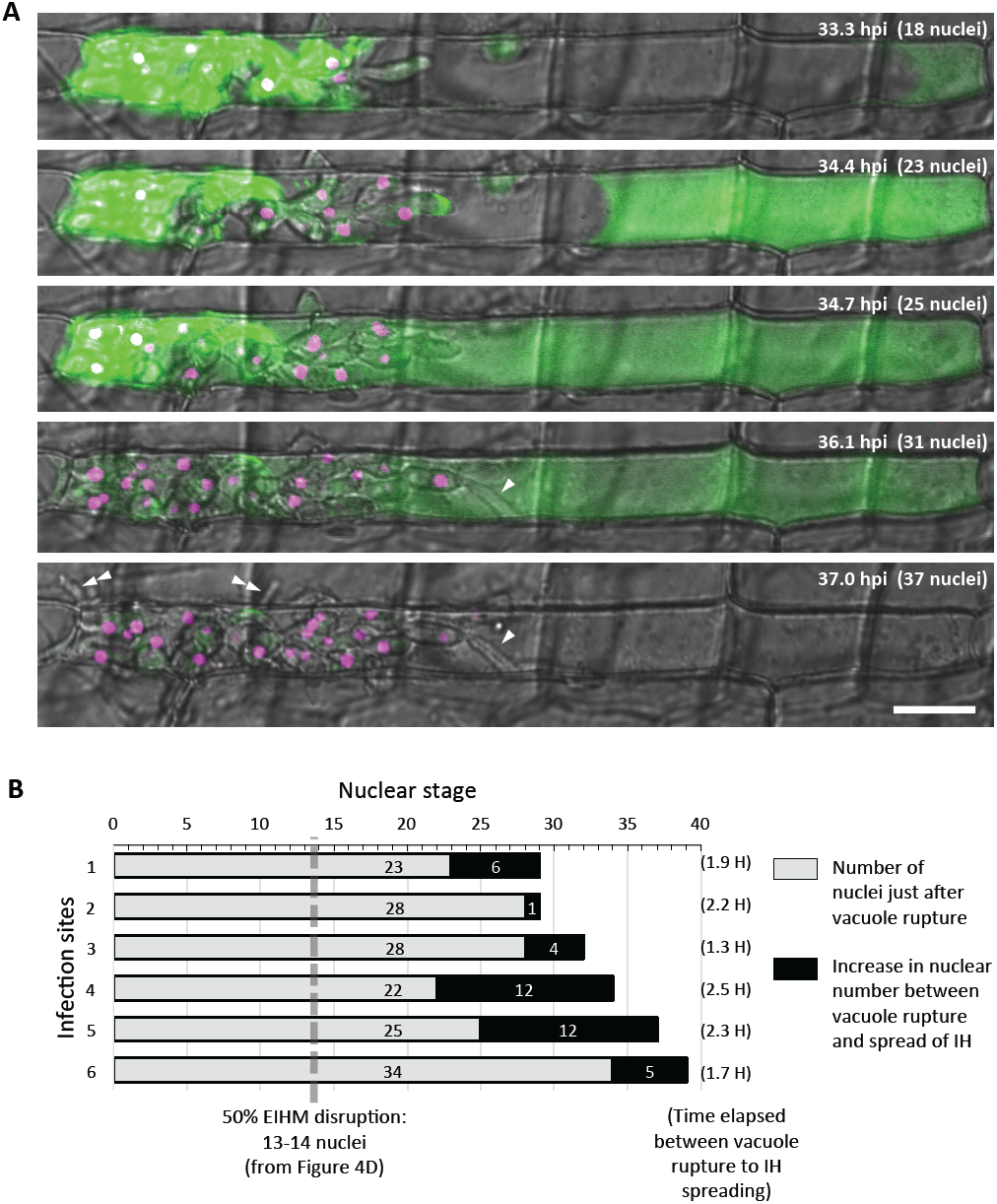
IH morphology changes after host cell death. (A) Representative time-lapse of *M. oryzae* CKF2187 expressing sec-GFP (green) and nuclear tdTomato (magenta) invading a rice cell from 33 to 37 hpi. Shown are single plane confocal images of merged bright-field and fluorescence. Nuclear stage is indicated in the upper right-hand corner together with hpi. Growth of IH become more filamentous after disruption of the vacuole (white arrowheads). The first IH to cross into the next host cell (double white arrowheads) originated from IH that had grown to be densely packed against the host cell wall before vacuole rupture. Bar = 20 μm. (B) Graphical summary showing six time-lapsed CKF2187 infections ranging from 32 to 40 hpi. Shown are the nuclear stages when vacuole rupture was observed (gray bars) and the relative increase in nuclear stage when IH were observed to spread into neighboring host cells (black bars). The time elapsed between vacuole rupture and IH spreading is shown in parenthesis, corresponding to the black bars. For additional context, the nuclear stage at which 50% EIHM disruption occurred (13-14 nuclei; Figure 4D; n = 390) is denoted by the dotted gray line.

Intriguingly, the growth of IH within the dead rice cell appeared morphologically distinct from bulbous IH; transitioning to growth that was more filamentous than the typical bulbous IH from which it arose (Figure 6A; white arrowheads). This transition was consistently observed throughout this study (Figure 2C and 5C). In addition, we noticed that the first IH to enter an adjacent rice cell were from IH which had been closely associated with the rice cell wall before the vacuole ruptured (Figure 6A; white double arrowheads). Despite the proximity of these IH to the cell wall crossing point, they did not invade adjacent cells for over an hour after rupture of the vacuole. This suggested that IH do not cross the rice cell wall into adjacent cells before the first-invaded cell dies.

### Dynamics of effector localization during host cell invasion

We investigated changes of subcellular locations of *M. oryzae* effectors during IH growth in first-and second-invaded cells using time-lapse imaging of rice cells invaded with *M. oryzae* strain CKF1616. This strain expresses apoplastic effector Bas4 fused to EGFP (Bas4:EGFP) together with cytoplasmic effector Pwl2 fused to mCherry and a nuclear localization signal (Pwl2:mCherry:NLS) (Khang et al., 2010). We found that during the early invasion of the first-invaded cell, Bas4:EGFP localized exclusively in EIHMx around IH, whereas Pwl2:mCherry:NLS preferentially accumulated in the BIC, in the nucleus of the invaded cell, and also in nuclei of nearby uninvaded cells (Figure 7A), consistent with previous reports (Khang et al., 2010). As IH continued to grow, Bas4:EGFP moved from the EIHMx into the rice cytoplasm but did not appear to move into adjacent cells (Figures 7B and 7C), a feature consistently observed with sec-GFP (Figures 2, 4C, 5A, 5C, and 6A). The initial cell-to-cell movement of Pwl2:mCherry:NLS (44.5 kD) and the subsequent containment of the spilled Bas4:EGFP (36 kD) and sec-GFP (26.9 kD) in the first-invaded cell suggest that plasmodesmata permeability changes from open to closed with EIHM disruption. The BIC-localization of Pwl2:mCherry:NLS and EIHMx-localization of the Bas4:EGFP appeared in second-invaded cells (Figure 7D), a typical pattern observed in compatible interactions (Mosquera et al., 2009; Khang et al., 2010).

**Figure 7.**
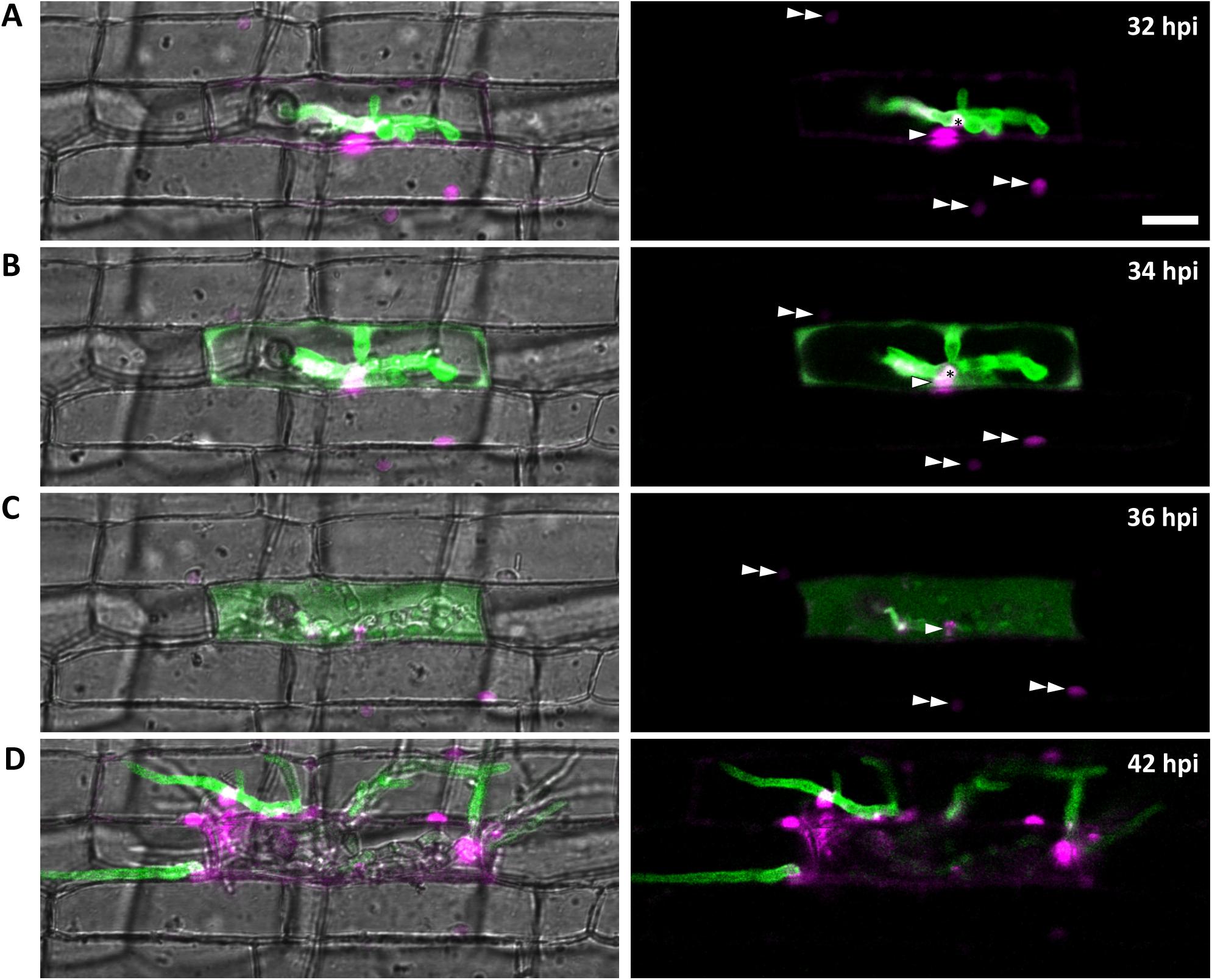
Effector localization changes during invasion of the first few host cells. *M. oryzae* CKF1616 expressing apoplastic effector Bas4:EGFP (green) along with cytoplasmic effector Pwl2:mCherry:NLS (magenta) invading rice. Shown are single plane merged fluorescence and bright-field (left panels), and merged fluorescence alone (right panels) confocal images of a time-lapse series from 32 to 42 hpi. Asterisk = BIC. Single white arrowhead = first-invaded host cell nucleus with Pwl2:mCherry:NLS fluorescence. Double white arrowhead = nuclei of uninvaded host cells with Pwl2:mCherry:NLS fluorescence. **(A)** During the early stage of invasion, the EIHM was still intact, causing Bas4:EGFP to be retained within the EIHMx. Pwl2:mCherry:NLS was localized at the BIC, in the nucleus of the invaded cell and in the nuclei of a few nearby cells. **(B)** The EIHM was disrupted, causing Bas4:EGFP to spill into the rice cytoplasm. **(C)** The vacuole ruptured, causing spilled Bas4:EGFP to homogenize throughout the host cell lumen. **(D)** IH invaded neighboring cells with Bas4:EGFP retained by new EIHMs. Pwl2:mCherry:NLS fluorescence increased upon invasion of neighboring host cells. By this time the first-invaded cell lacks significant levels of fluorescence. Bar = 20 μm.

## DISCUSSION

Plant-derived interfacial membranes are essential for the establishment and maintenance of biotrophy (Perfect and Green, 2001; Bozkurt et al., 2015), but little is known about the timing of interfacial membrane disruption, or what the consequences of this disruption are for both the pathogen and the infected host cell during hemibiotrophic invasion. In this study, we have provided evidence that the EIHM in first-invaded cells is disrupted in a manner dependent on IH growth stage and that EIHM disruption is an integral part of a successful infection. We developed a novel approach that allows EIHM integrity to be monitored during *M. oryzae* invasion of susceptible rice cells. This approach utilized sec-GFP that is localized in the EIHMx surrounded by the EIHM but spills into the host cytoplasm when the integrity of the EIHM is compromised (see examples in Figure 2). Using quantitative live-cell imaging coupled with sec-GFP and additional fluorescent reporters, we showed that the occurrence of EIHM disruption in the first-invaded cells was positively correlated with IH growth stage, and over 50% of infected cells possessed a disrupted EIHM when IH had grown to consist of more than 13 nuclei (13 nuclear stage) (Figure 4D). This growth stage corresponded to less than half of the colonization of the first-invaded host cell, considering that IH moved into adjacent cells at later than 29 nuclear stage (Figure 6B). The localization of sec-GFP was also found to be useful to visualize changes in the host vacuole and the permeability of plasmodesmata that occurred after EIHM disruption (Figures 2, 4C, 5, and 6).

What are the mechanisms underlying EIHM biogenesis and subsequent disruption? The answers to these questions remain largely unknown, though it is likely that EIHM integrity is linked to EIHM biogenesis, and thus aberrant EIHM biogenesis would result in the loss of EIHM integrity. Kankanala et al (2007) proposed that the EIHM is built *de novo* by redeploying host membranes toward the nascent rice-*M. oryzae* interface based on their FM4-64 dye-loading studies showing the dynamic association of host membrane tubules and round vesicles near the expanding EIHM. Recent studies with transgenic rice expressing GFP fusions with plasma membrane-localized proteins, such as OsCERK1, EL5, and LTi6b, further showed that the EIHM is continuous with the rice plasma membrane but appears to be distinct from it. OsCERK1:GFP and EL5:GFP were typically present in the invaginated host plasma membrane surrounding young IH but were absent from the EIHM surrounding the mature bulbous IH, whereas GFP:LTi6b continuously outlined IH (Mentlak et al., 2012; Kouzai et al., 2014; Mochizuki et al., 2015). These studies suggest that EIHM biogenesis begins with invagination of the host plasma membrane to surround early IH growth and subsequently transitions to *de novo* construction when IH differentiate bulbous growth. The latter likely involves the modulation of host membrane dynamics similar to those observed during interface biogenesis in other host-pathogen interactions (Koh et al., 2005; Micali et al., 2011; Bozkurt et al., 2015; Deeks and Sánchez-Rodríguez, 2016; Inada et al., 2016). Although the membrane source(s) and trafficking mechanism to build the EIHM remain unknown in the rice-*M. oryzae* interaction, the exhaustion of the source material or the perturbation of the trafficking processes may lead to the failure of EIHM biogenesis and expansion to accommodate growing IH. This hypothesis is supported by our results indicating that the EIHM initially loses integrity at the tips of late-stage IH and that this integrity loss is irreparable (Figure 3). Alternatively, *M. oryzae* may use a strategy similar to intracellular bacterial pathogen *Listeria monocytogenes* and protozoan pathogen *Toxoplasma gondii* that initially reside within a vacuole but later produce pore-forming proteins, Listeriolysin O and perforin-like protein 1, respectively, to disrupt the vacuole and reach the host cytoplasm (Kafsack et al., 2009; Hamon et al., 2012). It remains to be determined whether late-stage IH tips secrete pore-forming proteins to disrupt the EIHM and if, upon reaching the host cytoplasm, they contribute to permeabilization of the host vacuole membrane leading to gradual shrinkage and rupture of the vacuole.

The results presented in this study suggest that EIHM disruption and host cell death are landmarks that demarcate three distinct phases of growth within the first-invaded rice cell (Figure 8). First, the early biotrophic phase maintains hallmarks typical of biotrophy, in which IH grow in living host cells while being surrounded by the intact EIHM. Second, the late biotrophic phase begins when the EIHM is disrupted, causing IH to grow with increasingly direct contact with the living host cytoplasm. During this phase the rice vacuole progressively shrinks and eventually disrupts, which coincides with death of the invaded cell marking the end of biotrophy. Third, the transient necrotrophic phase takes place within the dead host cell, ending when biotrophy is reestablished upon invasion of adjacent rice cells. During the transient necrotrophic phase, IH switch from bulbous to filamentous-like growth. This necrotrophy-like growth was transient. That is, IH grew within the dead host cell for ∼1.3 to 2.5 h (Figure 6), which is relatively brief compared with the ∼12 h IH spend colonizing the first-invaded cell (Kankanala et al., 2007). It remains to be determined what triggers IH to move into adjacent cells. We found that IH were often closely associated with host cell walls even before vacuole rupture (early/late biotrophic phases), and that these IH were often the first to enter adjacent rice cells (Figure 6). Interestingly, however, IH did not move into adjacent cells until at least more than an hour after vacuolar rupture and host cell death (transient necrotrophic phase) (Figures 6). This suggests that the transient necrotrophic phase is required for IH cell-to-cell movement. Taken together, we propose that *M. oryzae* undergoes three distinct hemibiotrophic phases in each newly invaded cell during symptomless early invasion, and this lifestyle is followed by a complete transition to necrotrophy associated with macroscopic lesion development that typically occurs a few days after inoculation.

**Figure 8.**
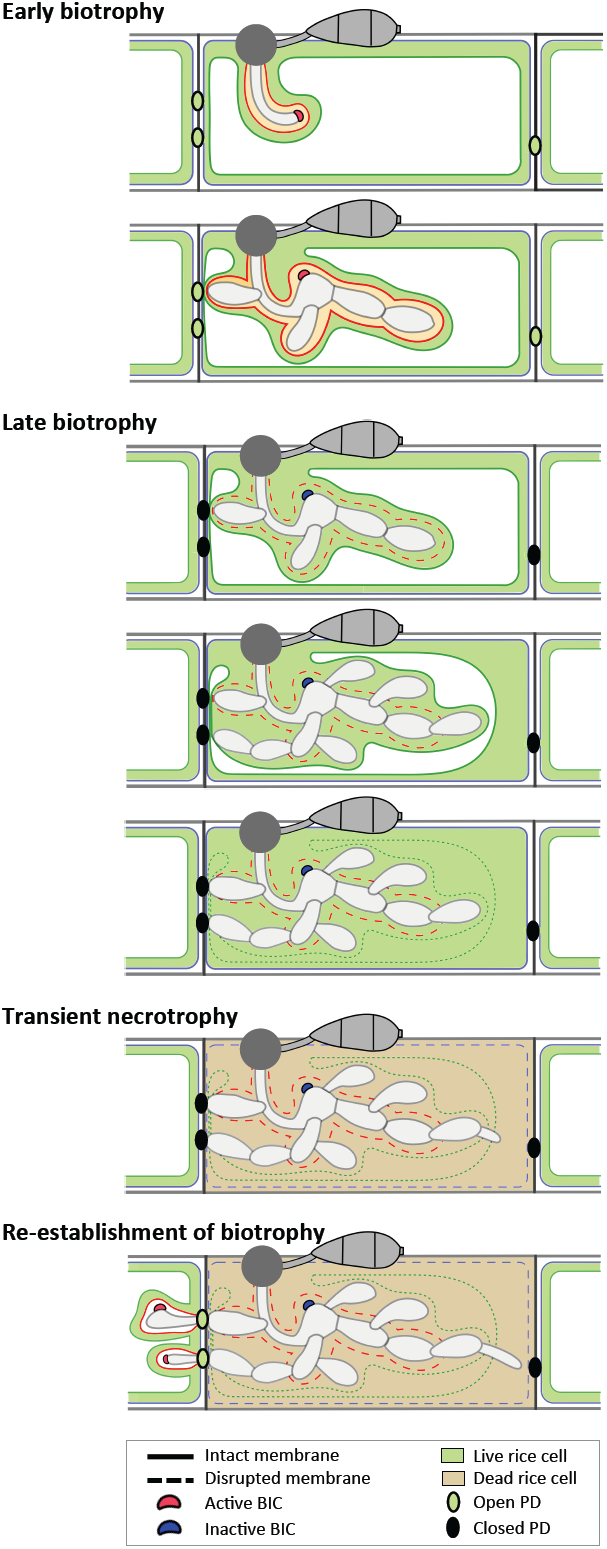
Model of the hemibiotrophic lifestyle of *M. oryzae* during invasion of the first and second rice cells. **Early biotrophy:** An initial invasion of a rice cell is achieved by a filamentous primary hypha, which differentiates into the first bulbous IH cell. The tip BIC positioned at the apex of the primary hypha is left at a subapical position when the first bulbous cell differentiates. Branched bulbous IH then arise from both the first bulbous cell and from the primary hypha. All IH are encased by an intact EIHM. The EIHM-encased IH also invaginate the vacuole, resulting in a thin layer of cytoplasm on the rice-facing side of the EIHM. Apoplastic effectors are retained within the EIHMx, while cytoplasmic effectors accumulate at the BIC, enter the host cytoplasm, and move symplastically through open plasmodesmata into adjacent cells. **Late biotrophy:** The EIHM disrupts, causing apoplastic effectors to spill from the EIHMx into the host cytoplasm and exposing IH to direct contact with the host cell cytoplasm. By this time, effector cell-to-cell movement has ceased due to closed plasmodesmata. The host vacuole progressively shrinks around growing IH, resulting in increased cytoplasmic volume. This eventually ends in rupture of the vacuole, causing the cytoplasm and vacuolar contents to homogenously mix. **Transient necrotrophy:** The plasma membrane becomes permeabilized when the vacuole ruptures, resulting in host cell death. This occurs in a contained manner without affecting the viability of adjacent host cells. Leading IH then differentiate more filamentous growth, which lasts at least over an hour before invasion of adjacent host cells. **Re-establishment of biotrophy:** The first IH to invade a neighboring cell often originates from IH which have grown to be in close association with the rice cell wall before vacuole rupture. Invasion of adjacent cells is biotrophic with formation of new BICs and EIHM as well as invagination of the vacuole. Cytoplasmic and apoplastic effectors are again delivered to the cytoplasm and EIHMx, respectively.

Our results showed that the viability loss of first-invaded rice cells coincided with the rupture of the central vacuole (Figure 5). Because the vacuole contains various hydrolytic enzymes, the rupture of the vacuole releases these enzymes into the cytoplasm where they degrade cellular organelles, eventually culminating in plant cell death (Jones, 2001; Hara-Nishimura and Hatsugai, 2011). The vacuole rupture is known to contribute to either disease resistance or susceptibility depending on pathogen lifestyle and the timing of the rupture relative to infection stage (Hatsugai et al., 2004; Hara-Nishimura and Hatsugai, 2011; Dickman and Fluhr, 2013; Mochizuki et al., 2015). Mochizuki et al (2015) used transgenic rice expressing vacuole membrane-localized GFP and showed that the vacuole gradually shrank and eventually ruptured in susceptible *M. oryzae*-invaded rice cells, consistent with our results. They further demonstrated that vacuole rupture caused critical damage to non-branched IH at the early infection stage but not to branched IH at the later infection stage, suggesting a fungal-driven mechanism for maintaining the integrity of the host vacuole until IH gain tolerance to vacuole rupture, a possibility that requires further investigation. Vacuole-mediated cell death in plants is regulated by vacuolar processing enzymes (VPEs) (Hatsugai et al., 2006; Hatsugai et al., 2015). In rice, five *VPE* (*OsVPE*) genes have been identified, and the expression levels *of OsVPE2* and *OsVPE3* were shown to increase during H_2_O_2_-induced vacuole rupture and cell death (Deng et al., 2011; Christoff et al., 2014). Our preliminary result showed that *OsVPE1* expression increased in *M. oryzae* infections at 32 hpi when most infected cells exhibited features of the late biotrophic phase (J. Zhu and C.H. Khang, unpublished results). We propose that death of first-invaded cells is vacuole-mediated and that *OsVPE* genes are involved in both *M. oryzae*-and abiotic stress-induced cell death in rice, although the role of different *OsVPE* genes may vary depending on the type of stress inflicted.

This study provides evidence that subcellular localization of *M. oryzae* effectors change during IH growth in first-invaded cells (Figure 7). During the early biotrophic phase, fluorescently-tagged effectors Bas4 (apoplastic effector) and Pwl2 (cytoplasmic effector) exhibited distinct localization patterns; Bas4 accumulated in EIHMx, whereas Pwl2 preferentially accumulated in BICs, entered the host cytoplasm, and moved into surrounding cells (Figure 7), consistent with previous results (Khang et al., 2010). Bas4 was then relocalized during the late biotrophic phase when the EIHM was disrupted; spilling from the EIHMx into the host cytoplasm. These results suggested that all *M. oryzae* effectors eventually enter the host cytoplasm either by translocation across the intact EIHM (cytoplasmic effectors) during the early biotrophic phase or by spilling through the disrupted EIHM (apoplastic effectors) during the late biotrophic phase. Apoplastic effector re-localization has significant implications in the study of effectors: (1) Effectors are generally classified into either cytoplasmic effectors or apoplastic effectors, depending on their localization in the host cytoplasm or in the apoplast (interfacial compartment or intercellular space), respectively (Schornack et al., 2009; Giraldo and Valent, 2013; Lo Presti and Kahmann, 2017). We suggest that these effector classifications, however, must be defined in the context of infection stages, at least for *M. oryzae* effectors such as Bas4 that are localized in the EIHM compartment but are subsequently re-localized in the host cytoplasm; (2) Live-cell imaging of *M. oryzae* expressing fluoresecently-tagged effectors has been instrumental for determining the identity of an effector as apoplastic or cytoplasmic and for investigating the mechanism by which cytoplasmic effectors are translocated into the host cytoplasm (Mosquera et al., 2009; Khang et al., 2010; Mentlak et al., 2012; Park et al., 2012; Ribot et al., 2013; Nishimura et al., 2016; Sharpee et al., 2017). This approach requires that individual infection sites be assessed for EIHM integrity, such as use of sec-GFP, to differentiate effector translocation across the intact EIHM from effector entry through the disrupted EIHM as suggested by earlier studies (Mosquera et al., 2009; Khang et al., 2010; Giraldo and Valent, 2013); (3) Given the intimate link between effector localization and function (Giraldo and Valent, 2013; Sharpee et al., 2017), *M. oryzae* apoplastic effectors may play roles in both the apoplast and the cytoplasm, depending on their phase-specific localizations. Three *M. oryzae* apoplastic effectors, Slp1, Bas4, and Bas113, have been shown to localize in the EIHMx, and Slp1 was further determined to function as a LysM protein that sequesters chitin to suppress host immunity (Mosquera et al., 2009; Mentlak et al., 2012; Giraldo et al., 2013). It is an exciting possibility that these and many yet-to-be-identified apoplastic effectors may have host targets in both the apoplast and the cytoplasm.

Although the mechanism of how effectors are translocated across the interfacial membrane remains unknown for most filamentous pathogens, increasing evidence suggests that BICs formed in *M. oryzae*-invaded rice cells function as the site of translocation into the rice cytoplasm across the intact EIHM (Khang et al., 2010; Giraldo et al., 2013). We propose that BICs undergo three developmental and functional stages in first-invaded cells: in the first stage, a single ‘tip-BIC’ appears at the tip of the filamentous primary hypha; in the second stage, the tip BIC becomes the ‘early side-BIC’ when the filamentous hypha differentiates bulbous growth; in the third stage, the early side-BIC remains on the side of the first bulbous IH cell as the ‘late side-BIC’ while IH continue to proliferate in the rice cell (Figure 8). Host cytoplasmic dynamics appear to be focused in the vicinity of the tip BIC and the early side-BIC, and these BICs strongly accumulate fluorescently-tagged cytoplasmic effectors, whereas the late side-BIC shows weaker intensity of effector-associated fluorescence (Khang et al., 2010). We hypothesize that the tip-and the early side-BICs are actively performing their presumed function in effector delivery during the early biotrophic phase when the EIHM is intact, and the late side-BIC is a remnant that has ceased to deliver effectors. This is consistent with our preliminary evidence that expression of the BIC-localized cytoplasmic effector gene *PWL2* is strongly induced at early infection stages when the tip-and early side-BIC were present (unpublished). A future research tool that can directly demonstrate the role of BICs in effector translocation is needed to test this hypothesis. It has been an intriguing question why there is only a single BIC in each first-invaded cell. It may be because the BIC is required for the delivery of cytoplasmic effectors into host cells only during the early biotrophic phase while the EIHM is intact; becoming obsolete once the EIHM is disrupted.

Previous studies suggest that *M. oryzae* exploits open plasmodesmata for cell-to-cell movement of effectors and of IH during biotrophic invasion (Kankanala et al., 2007; Khang et al., 2010). Our results expand these studies and further suggest that plasmodesmata permeability changes in a manner specific to infection phase. Evidence for the plasmodesmata being open during the early biotrophic phase comes from Pwl2:mCherry:NLS (44.5 kD), which entered the host cytoplasm across the intact EIHM and moved into surrounding cells (Figure 7A). Evidence for the plasmodesmata closure in the late biotrophic and transient necrotrophic phases comes from Bas4:EGFP (36 kD) and sec-GFP (26.9 kD), both of which entered the invaded cell through the disrupted EIHM and accumulated there without diffusion into adjacent cells (Figure 7B and 7C). We propose that infection phase-dependent plasmodesmata dynamics are integral to *M. oryzae*’s successive biotrophic invasion. That is, during the early biotrophic phase, plasmodesmata serve as a conduit for cytoplasmic effectors to move into surrounding uninvaded rice cells where they are presumed to prepare rice cells for invasion (Khang et al., 2010). During the subsequent infection phases when the viability of the first-invaded cell declines and is eventually lost, plasmodesmata become closed, which prevents death signals from spreading into uninvaded cells, thus keeping these cells unaffected and viable. Subsequently, plasmodesmata are exploited by IH to move into adjacent viable host cells (Kankanala et al., 2007).

How plasmodesmata permeability is regulated during rice blast disease remains an open question. It is generally known that pathogen infections induce plasmodesmata closure by recruiting plasmodesmata-associated molecules such as callose (a β-1,3 glucan polymer) and that such plasmodesmata closure is linked to host immunity (Lee, 2015). In particular, recent studies in *Arabidopsis thaliana* suggest that recognition of chitin (pathogen-associated molecular pattern, PAMP, from fungal pathogens) by the chitin pattern recognition receptor LYM2 (also known as AtCEBiP) leads to plasmodesmata closure (Faulkner et al., 2013) and also that the plasmodesmata-localized protein 5 mediates callose deposition at plasmodesmata in a manner depending on the defense hormone salicylic acid (Lee and Lu, 2011). It is an interesting possibility that PAMP-triggered plasmodesmata closure in rice is suppressed when IH grow within the EIHM (the early biotrophic phase) but is then activated when IH are exposed to the host cytoplasm after EIHM disruption, which might result in increased PAMP recognition by rice PRRs (the late biotrophic phase). Mentlak et al (2012) showed that *M. oryzae* secretes the apoplastic effector Slp1 to sequester chitin released from IH growing within the EIHM and thus prevents chitin from being recognized by the rice chitin PRR CEBiP. It remains to be determined whether *M. oryzae* Slp1 and rice PRRs, including CEBiP, play a role in plasmodesmata regulation during rice blast disease. Although the precise mechanism of how IH cross the cell wall to invade adjacent cells after the transient necrotrophic phase remains unknown, it may involve modulation of closed plasmodesmata, for example degrading plasmodesmata callose using hydrolytic enzymes such as β-1, 3–glucanases and β-glucosidases to reopen them. Understanding how plasmodesmata permeability is regulated during *M. oryzae* invasion and how the plasmodesmata dynamics is linked to host susceptibility and resistance will offer potential targets that can be exploited to control blast disease.

## METHODS

### Strains, Fungal Transformation and Plasmid Construction

*M. oryzae* wild-type strain O-137, isolated from rice (*Oryza sativa*) in China (Orbach et al., 2000), was used as a recipient strain to generate fungal transformants using *Agrobacterium tumefaciens*-mediated transformation (Khang et al., 2006). We used the rice strain YT-16 highly susceptible to *M. oryzae* O-137 (Kankanala et al., 2007)and all O-137-derived transformants used in this study. See Supplemental Table 1 online for the list of *M. oryzae* transformants. The Dendra2 gene was PCR-amplified from tol2-mpx-Dendra2 (a gift from Anna Huttenlocher; Addgene plasmid # 29574; (Yoo and Huttenlocher, 2011)) using the primers CKP303: 5’-GGATCCATGAACACCCCGGGAATTAAC-3’ and CKP304: 5’-TGTACAGCCACACCTGGCTGGG-3’, underlined for *Bam*HI and *Bsr*GI sites, respectively. The *BAS4* promoter and its entire 102-amino acid coding sequence (1.3 kb *Eo*RI-*Bam*HI fragment (Khang et al., 2010) and Dendra2 were cloned together with the Nos terminator (0.3 kb *Bsr*GI-*Sal*I fragment from pBV360 (same as pAN583; Nelson et al., 2007) in the binary vector pBGt to generate pCK1244 (*BAS4*:Dendra2:Terminator). The *BAS4* promoter and signal peptide-encoding sequence were cloned together with the *EGFP* and the *Neurospora crassa* β-tubulin gene terminator in the binary vector pBHt2 to generate pBV324 (sec-GFP construct) (Khang et al., 2010)). The *M. oryzae* ribosomal protein 27 gene (P27) promoter was used to construct the constitutive expression plasmid pCK1292 for cytoplasmic tdTomato (Jones et al., 2016b). The EGFP gene was obtained from Clontech, and the tdTomato was isolated from pAN582 (Nelson et al., 2007). The P27 promoter and histone H1 gene from *N. crassa*, which was isolated from pBV229 (Shipman et al., 2017), was cloned together with tdTomato at the upstream of the sec-GFP construct in pBV324 to generate pCK1312. See Supplemental Table 2 online for the list of plasmids used.

### Infection Assays

Rice sheath inoculations were performed as previously described (Kankanala et al., 2007). Briefly, excised leaf sheaths (5-9 cm long) from 2 to 3 weeks old plants were inoculated by injecting a spore suspension (5 × 10^4^ spores/ml in sterile water) into the hollow interior of the sheath. The inner epidermal layer of the inoculated sheath was hand-trimmed for confocal microscopy.

### Staining and Plasmolysis

Propidium iodide (PI) was prepared to a 10 μg/ml working solution by diluting 10 μl of stock solution (catalog No. P3566; 10 ml of 1 mg/ml solution in water; ThermoFisher) in 990 μl of water. Trimmed leaf sheaths were submerged in the PI working solution for 15 minutes and then mounted in the same solution for microscopy. FM4-64 was prepared to a 17 mM aqueous stock solution by adding 9.2 μl of sterile distilled water to 100 μg of FM4-64 powder (catalog No. T13320; 10 x 100 μg; ThermoFisher) and stored at −20 °C. Trimmed leaf sheaths were incubated in a 17 mM aqueous working solution for 1 hour, washed with water, and then incubated for four more hours prior to microscopy. Fluorescein diacetate (FDA; catalog No. F7378, 5 g powder; Sigma) was dissolved in acetone to make a stock concentration of 1 mg/ml. A working solution of FDA (2 μg/ml, 0.2% acetone) was prepared by diluting 2 μl of the stock solution in 1ml of water. Sucrose-induced plasmolysis was performed by replacing the mounting solution of water with a 0.5 M sucrose solution and incubated for 25 minutes before microscopy.

### Confocal Microscopy

Confocal microscopy was performed on a Zeiss Axio Imager Z1 inverted microscope equipped with a Zeiss LSM 710 system using Plan-Apochromat 20×/0.8 NA and Plan-Neofluor 40×/1.3 NA (oil) objectives. Excitation/emission wavelengths were 488 nm/496 to 544 nm for GFP and fluorescein, 543 nm/565 to 617 nm for mCherry and tdTomato, 543 nm/580 to 640 nm for PI, and 543 nm/613 to 758 nm for FM4-64. Images were acquired using the Zen Black 2011 software. Images were processed using the Zen Black software (version 10.0, Zeiss). For long interval time-lapse imaging, the coverslip was removed and water was added to the slide in between images to prevent dehydration and to allow gas exchange to occur. Selective photoconversion of Bas4:Dendra2 was performed by irradiating a region of interest with the 405 nm laser line (100 % output power and a pixel dwell time of 1.58 μs with 250 iterations) using the 40x objective lens at a zoom factor of 2. Excitation/emission wavelengths for imaging unconverted green Dendra2 were 488 nm/496 to 554 nm and 543 nm/560 to 675 nm for imaging converted red Dendra2.

### Supplemental data

The following materials are available in the online version of this article.

**Supplemental Figure 1.** FM4-64 labels IH septa only when sec-GFP is spilled into the host cell.

**Supplemental Figure 2.** Sec-GFP spills into the rice cytoplasm after EIHM disruption. **Supplemental Figure 3.** Image analysis – Brightness and contrast adjustment to reveal instances of low-intensity sec-GFP fluorescence in the host cell.

**Supplemental Figure 4.** Variation in host-localized sec-GFP fluorescence patterns.

**Supplemental Table 1.** *Magnaporthe oryzae* transformants used in this study.

**Supplemental Table 2.** Key Plasmids Used in This Study

## Acknowledgements

We thank all members of the Khang Lab (http://www.khanglab.org/), especially current member Mariel Pfeifer, for their help and discussion. We acknowledge the assistance of the Biomedical Microscopy Core at the University of Georgia with imaging using a Zeiss LSM 710 confocal microscope. This work was supported by the Agriculture and Food Research Initiative competitive grants program, Award number 2014-67013-21717 from the USDA National Institute of Food and Agriculture.

## Authors’ contributions

CHK conceived and designed the experiments. KJ, JZ, CBJ and DWK performed the experiments. KJ, JZ, CBJ, DWK and CHK analyzed the data and wrote the paper.

## Supplemental Figure Legends

**Supplemental Figure 1.**
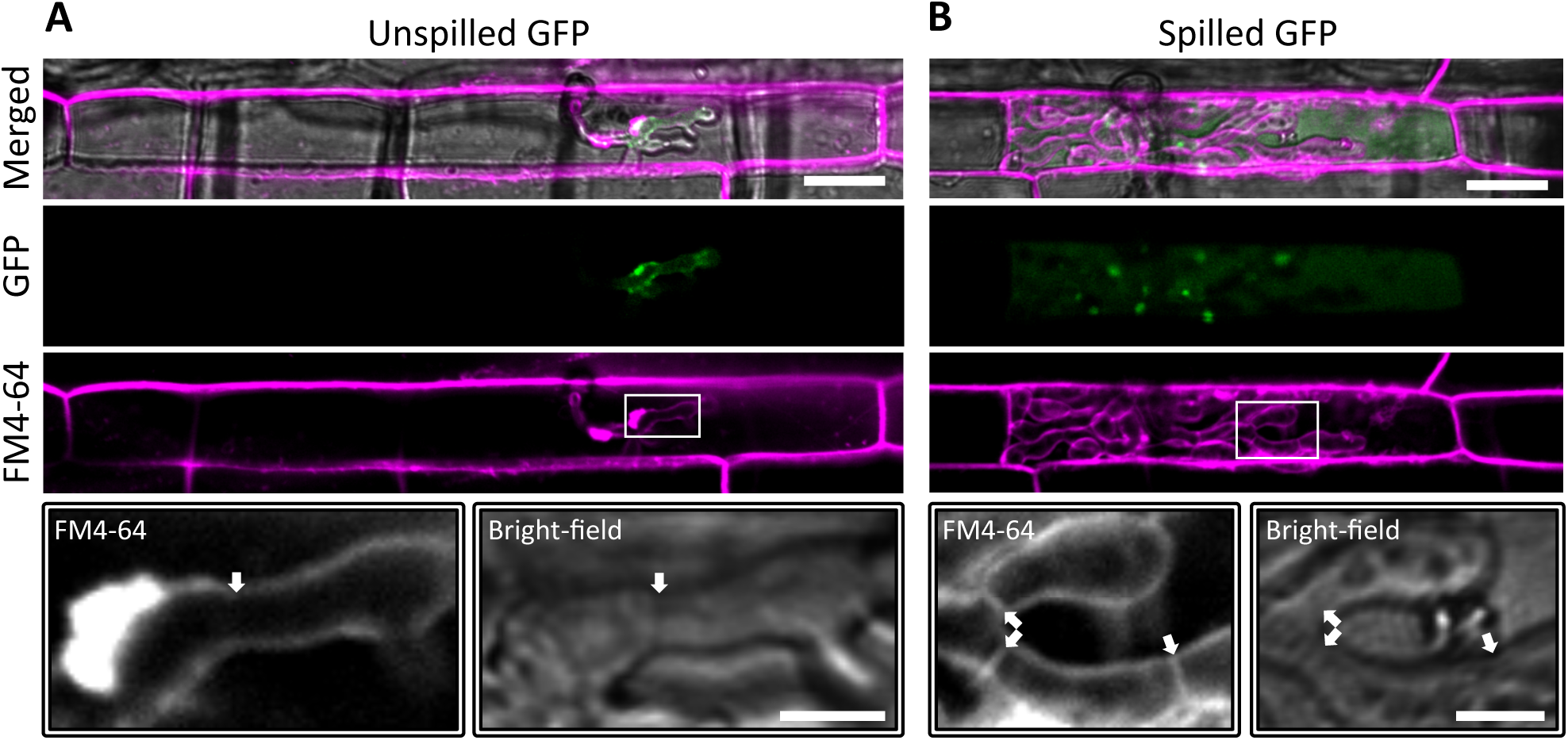
FM4-64 labels IH septa only when sec-GFP is spilled into the host cell. **(A)** and **(B)** *M. oryzae* strain CKF2180 (sec-GFP; green) invading rice cells. Inoculated sheaths were pulse-stained with FM4-64 (shown in magenta) for one hour, washed with water, and then incubated for four hours prior to microscopy. Shown are single plane merged fluorescence, split fluorescence, and bright-field confocal images. Bars = 20 μm (full size images) and 5 μm (insets). **(A)** An infection at 28 hpi shows sec-GFP exclusively outlining IH. Inset shows a region of IH enlarged to demonstrate the absence of FM4-64 labelling (pseudo-colored white) near the septum (white arrow). Both EIHMx-localized sec-GFP and the absence of FM4-64 labelling from IH septa were consistent with an intact EIHM preventing the diffusion of either fluorophore. **(B)** A different infection at 32 hpi shows sec-GFP spilled into the host cell. Inset shows a region of IH enlarged to show positive FM4-64 labelling of fungal membranes at three septa (white arrows). Both the host-localized sec-GFP and fungal labelling of FM4-64 were consistent with a disrupted EIHM.

**Supplemental Figure 2.**
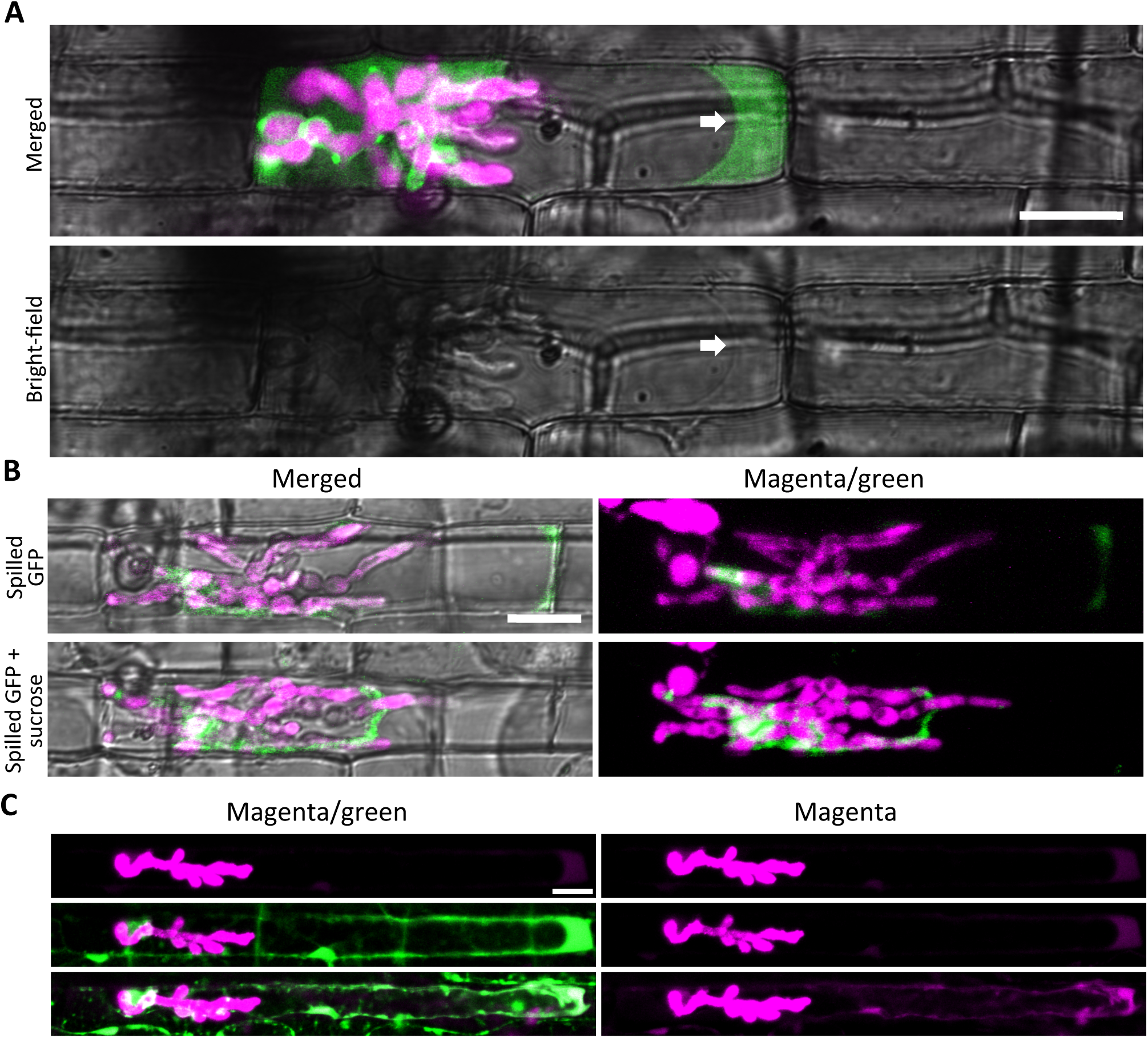
Sec-GFP spills into the rice cytoplasm after EIHM disruption. **(A)** and **(B)** *M. oryzae* CKF1996 expressing sec-GFP (green) and cytoplasmic tdTomato (magenta) invading rice. Shown are single plane confocal images of both merged fluorescence and bright-field (**A**, top; **B**, left), or bright-field alone (**A**, bottom), and a merged fluorescence projection of 15 z-slices with 2 μm each (**B**, right). Bars = 20 μm. (**A**) Infection at 30 hpi with sec-GFP spilled into the host cell, indicating the EIHM was disrupted. The vacuole membrane is visible in the bright-field (white arrows). (**B**) Infection at 32 hpi with sec-GFP in the host cell. After the top image was taken water was replaced with 0.5 M sucrose to induce plasmolysis. After 25 minutes, host-localized sec-GFP was retracted from the cell wall and remained excluded from the vacuole, demonstrating that sec-GFP was indeed localized within the host cytoplasm. Note that IH shifted slightly after the host cell was plasmolyzed. (**C**) *M. oryzae* CKF3267 expressing secreted mCherry (magenta) invading a rice cell between 31 and 33 hpi. Shown are single plane merged or split fluorescence confocal images of the same infection site. Like sec-GFP, secreted mCherry spilled into the host cytoplasm (top), indicating a disrupted EIHM. The rice sheath was then stained with 0.2 μg/ml fluorescein diacetate (FDA). FDA is converted to its fluorescent form in the rice cytoplasm where it is then retained (Jones et al., 2016b). FDA fluorescence (green) co-localized with secreted mCherry in the host cell, confirming cytoplasmic localization of spilled mCherry (middle). This was further confirmed by subsequently inducing plasmolysis with 0.5 M sucrose, which caused retraction of the colocalized spilled mCherry and FDA fluorescence from the cell wall as expected (bottom). The time elapsed between each image was 30 minutes. Bar = 20 μm.

**Supplemental Figure 3.**
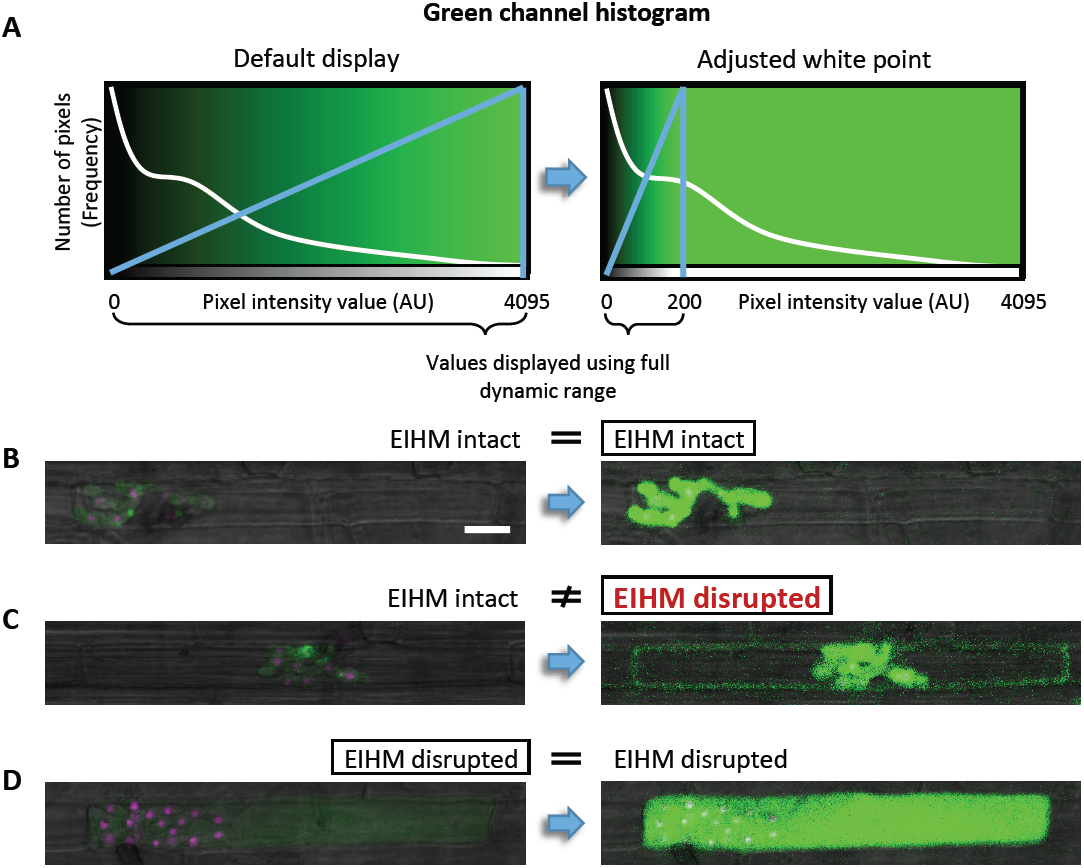
Image analysis – Brightness and contrast adjustment to reveal instances of low-intensity sec-GFP fluorescence in the host cell. During the initial stages of our quantitative analysis of sec-GFP localization and nuclear stage, we discovered that some infected host cells contained low-intensity green fluorescence in the cytoplasm that was impossible or nearly impossible to detect at the default display setting. Infections showing this pattern were prone to being misinterpreted as possessing an intact EIHM unless a more detailed image analysis was performed. To maximize the sensitivity of our EIHM integrity assay we empirically derived an appropriate adjustment to the image display settings in the Zen software (black edition) that consistently revealed instances of low intensity green fluorescence in the host cytoplasm (Fig. 4C and 4D; n = 390). Ultimately we found that sufficient brightness and contrast for resolving background noise and low-intensity green fluorescence was achieved by adjusting the white point in the green fluorescence channel histogram from the default maximum of 4095 (for 12-bit images) **(A;** left) to 200 **(A;** right). This produced a display of the image where pixel intensity values 0-200 were proportionally increased in intensity in order to populate the full dynamic range (grayscale), while values 201-4096 were displayed as saturated. Once the new white point was applied to an image, individual z-stacks were inspected for presence of host-localized green fluorescence at low-intensity. Shown in **(B)** through **(D)** are single plane merged bright-field and fluorescence confocal images of *M. oryzae* CKF2187 infections expressing sec-GFP (green) and H1:tdTomato (magenta) during invasion of the first rice cell between 29 and 31 hpi. Each shows a representative outcome of the image analysis. At default display settings, infection **(B)** appeared to have EIHMx-exclusive sec-GFP localization, indicating an intact EIHM. After the white point was lowered in the green channel, the sec-GFP localization pattern was confirmed as EIHMx-exclusive. Similar to **(B)**, infection **(C)** initially appeared to have an intact EIHM at the default display setting. However, after reducing the white point, low-intensity sec-GFP fluorescence was revealed to be present in the host cytoplasm, thus reversing the initial scoring and highlighting the importance of careful image analysis. Infection **(D)** was readily discernable as having a disrupted EIHM with both the default and adjusted display settings. Note that infections with this pattern (host-localized sec-GFP, disrupted vacuole) were usually identifiable with the default display. Bar = 20 μm, scale is equivalent for all images.

**Supplemental Figure 4.**
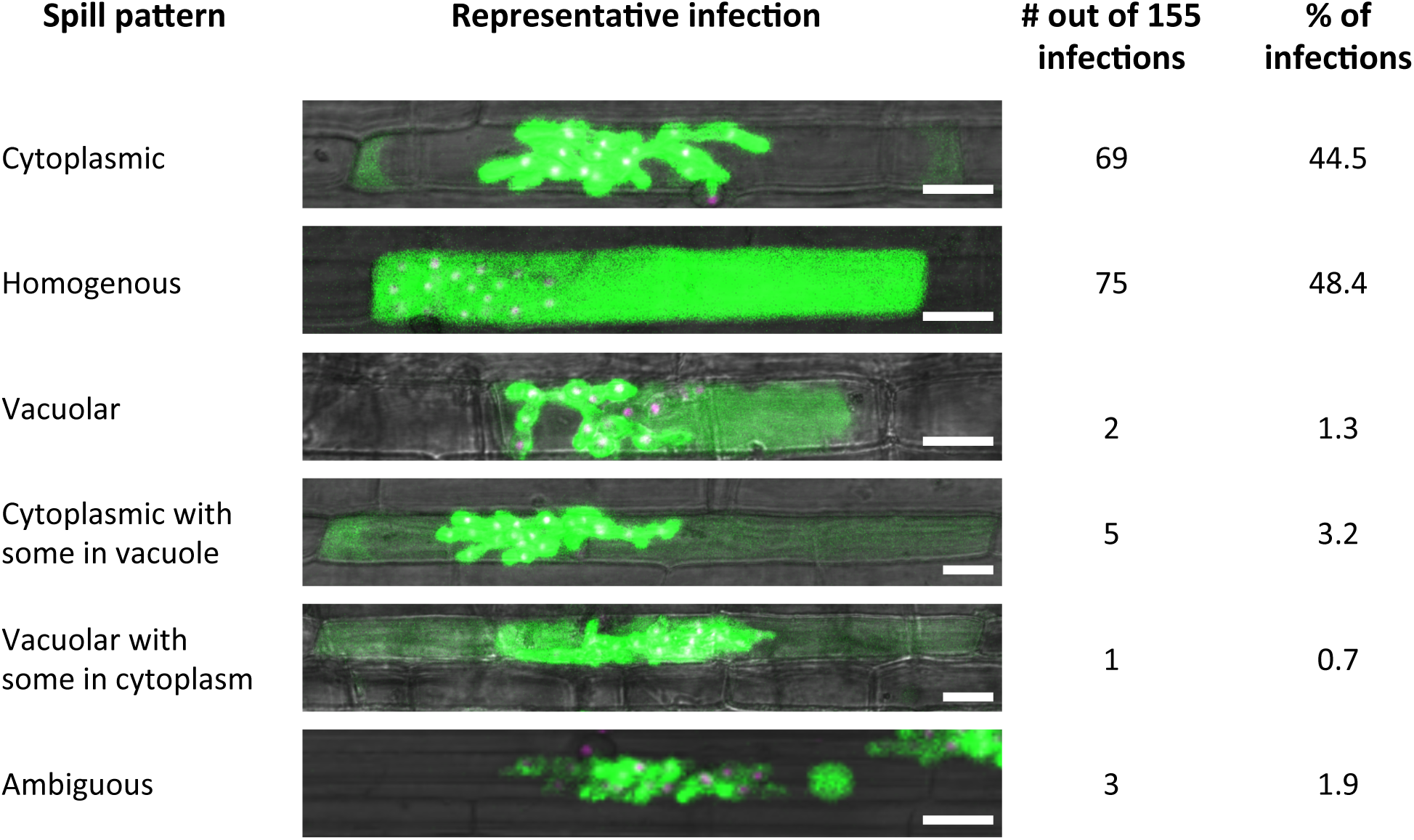
Variation in host-localized sec-GFP fluorescence patterns. Our quantitative analysis of sec-GFP (green) localization in the context of nuclear stage (magenta) for *M. oryzae* CKF2187 infections between 28 and 33 hpi revealed 155 infections (out of 390) with host-localized sec-GFP (Figure 4 D). The majority of these patterns were cytoplasmic (44.5 %) or homogenous throughout the rice cell (48.4 %) with the remaining 7.1% showing sec-GFP fluorescence: (1) inside only the vacuole (1.3%), (2) in both the cytoplasm and vacuole with higher intensity in the cytoplasm (3.2%), (3) in both the cytoplasm and vacuole with higher intensity in the vacuole (0.7 %), and (4) ambiguous host-localization (1.9%). Together, these data indicated that spilled sec-GFP was typically found to be cytoplasmic or homogeneous, however, it could occasionally spill into the vacuole, or other combinations of host compartments. Shown are single plane merged fluorescence and bright-field confocal images of representative CKF2187 infections for each variation of host-localized sec-GFP fluorescence. Bars = 20 μm.

---

**Supplemental Table 1.** *Magnaporthe oryzae* transformants used in this study.

| Strain | Used in | Description<br>[Recipient strain; Plasmid used] |
| --- | --- | --- |
| CKF315<sup>a</sup> | Figures 5A and 5C | Transformant expressing the Bas4 signal peptide fused to EGFP under control of the *BAS4* promoter [O-137<sup>b</sup>; pBV324]; same as KV97 (Mosquera et al., 2009) |
| CKF1616 | Figure 7 | Transformant expressing both a fusion of the Pwl2 coding sequence with mCherry:NLS under control of the *PWL2* promoter and a fusion of the Bas4 coding sequence with EGFP under control of the *BAS4* promoter [O-137; pBV591]; same as KV121 (Khang et al., 2010) |
| CKF1737 | Figure 1B | Transformant expressing a fusion of the Bas4 coding sequence with Dendra2 under control of the *BAS4* promoter [O-137; pCK1244] |
| CKF1996 | Figures 2B, 2C, and 3A;<br>Supplemental Figure 2A and 2B | Transformant expressing both a fusion of the Bas4 signal peptide fused to EGFP under control of the *BAS4* promoter and a cytoplasmic tdTomato under control of the constitutive promoter of the *M. oryzae* ribosomal protein 27 [CKF315; pCK1292] |
| CKF2180<sup>a</sup> | Supplemental Figure 1 | Transformant expressing the Bas4 signal peptide fused to EGFP under control of the *BAS4* promoter [O-137; pBV324] |
| CKF2187 | Figures 4B, 4C, and 6A;<br>Supplemental Figures 3B, 3C, 3D, and 4 | Transformant expressing both a fusion of the Bas4 signal peptide fused to EGFP under control of the *BAS4* promoter and a fusion of histone H1 with tdTomato under control of the constitutive promoter P27 [O-137; pCK1312] |
| CKF3267 | Supplemental Figure 2C | Transformant expressing the Bas4 signal peptide fused to mCherry under control of the *BAS4* promoter [O-137; pCK1594] |

<sup>a</sup>Two independent transformants containing the same construct showing consistent fluorescence patterns
<sup>b</sup>*M. oryzae* strain O-137 was isolated from rice (*Oryza sativa*) in China (Orbach, et al. 2000).

**Supplemental Table 2.** Key Plasmids Used in This Study.

| Clone | Description |
| --- | --- |
| pBV324 | 1 kb *EcoRI*-*Bam*HI fragment containing the *BAS4* promoter and 72 bp *Bam*HI-*Pst*I fragment encoding the Bas4 signal peptide cloned in the *Eco*RI and *Pst*I sites of pBV176 (C-terminal translational fusion of EGFP to the Bas4 signal peptide); Khang et al., 2010 |
| pBV591 | P<sub>PWL2</sub>:PWL2CDS:mCherry:NLS:Nopaline synthase gene (Nos) terminator and P<sub>BAS4</sub>:BAS4CDS:EGFP:Ter cloned in *Eco*RI and *Hind*III sites of pBHt2; Khang et al., 2010 |
| pCK1244 | 1.3 kb *Eco*RI-*Bam*HI fragment containing the *BAS4* promoter and coding sequence from pBV587, 0.7 kb *Bam*HI-*Bsr*GI fragment containing Dendra2 from pCK1206, and 0.3 kb *Bsr*GI-*Sal*I fragment containing the Nos terminator from pBV360 (same as pAN583; Nelson et al., 2007) cloned into *Eco*RI-*Sal*I sites of pBHt2 |
| pCK1292 | 0.5 kb *Eco*RI-*Bam*HI fragment containing the P27 promoter from pBV167 and 1.7 kb *Bam*HI-*Hind*III fragment containing tdTomato from pBV359 (same as pAN582; Nelson et al., 2007) cloned into *Eco*RI-*Hind*III sites of pBGt |
| pCK1312 | 2.2 kb *Eco*RI-*Bam*HI fragment of pBV229 containing the P27 promoter and histone H1 and 1.7 kb *Bam*HI-*Sal*I fragment of pBV359 containing tdTomato cloned into *Eco*RI-*Xho*I sites of pBV324 |
| pCK1594 | 1 kb *Eco*RI-*Bam*HI fragment containing the *BAS4* promoter and sequence encoding the signal peptide from pBV722 and 1 kb *Bam*HI-*Hind*III fragment of containing mCherry:Nos terminator cloned into *Eco*RI-*Hind*III sites of pBGt |

